# Base editor scanning charts the DNMT3A activity landscape

**DOI:** 10.1101/2022.04.12.487946

**Authors:** Nicholas Z. Lue, Emma M. Garcia, Kevin C. Ngan, Ceejay Lee, John G. Doench, Brian B. Liau

**Affiliations:** Department of Chemistry and Chemical Biology, Harvard University, Cambridge, MA 02138, United States; Broad Institute of Harvard and MIT, Cambridge, MA 02142, United States

## Abstract

DNA methylation is critical for regulating gene expression, necessitating its accurate placement by enzymes such as the DNA methyltransferase DNMT3A. Dysregulation of this process is known to cause aberrant development and oncogenesis, yet how DNMT3A is regulated holistically by its three domains remains challenging to study. Here we integrate base editing with a DNA methylation reporter to perform in situ mutational scanning of *DNMT3A* in cells. We identify mutations throughout the protein that perturb function, including ones at an interdomain interface that block allosteric activation. Unexpectedly, we also find mutations in the PWWP domain, a histone reader, that modulate enzyme activity despite preserving histone recognition and protein stability. These effects arise from altered PWWP domain DNA affinity, which we show is a noncanonical function required for full activity in cells. Our findings highlight mechanisms of interdomain crosstalk and demonstrate a generalizable strategy to probe sequence-activity relationships of nonessential chromatin regulators.

## Introduction

At every stage of mammalian life, chromatin regulation is essential for directing and safeguarding physiological processes. DNA methylation is a key chromatin modification that occurs throughout the human genome, primarily at CpG dinucleotide sites^1^. It has diverse functions, but canonically silences gene expression when present at promoter CpG islands^2^. DNA methylation is installed by the de novo DNA methyltransferases (DNMTs) DNMT3A and DNMT3B, which have long been recognized as essential for mammalian development^1,3^. Indeed, mutations in *DNMT3A* lead to developmental disorders, such as Tatton-Brown-Rahman Syndrome (TBRS)^4^ and Heyn-Sproul-Jackson syndrome (HESJAS)^5^. Moreover, *DNMT3A* mutations are important drivers of clonal hematopoiesis^6^ and acute myeloid leukemia (AML)^7,8^, as loss of DNMT3A function confers a proliferative advantage to hematopoietic stem cells, extending their renewal in vivo and promoting leukemic transformation^9–11^.

DNMT3A function is regulated by complex mechanisms involving its three domains. Beyond its direct role in catalysis, the methyltransferase (MTase) domain harbors interfaces enabling DNMT3A to complex with other copies of itself, DNMT3B, or the catalytically inactive homolog DNMT3L, thereby shaping its activity^8,12–16^. The two N-terminal domains, the ATRX-DNMT3-DNMT3L (ADD) and Pro-Trp-Trp-Pro (PWWP) domains, are histone reader modules recognizing histone H3 unmethylated at lysine 4 (H3K4me0) and dimethylated at lysine 36 (H3K36me2), respectively^17–19^. H3K4me0 recognition activates DNMT3A by repositioning its ADD domain, which otherwise blocks the DNA substrate binding site^17^. Although a similar mechanism involving the PWWP domain has not been elucidated, recent studies have found that H3K36me2 stimulates DNMT3A activity selectively over H3K36me0 in vitro^19,20^, raising the question of whether allosteric crosstalk between the PWWP and MTase domains might exist.

Understanding the interplay between DNMT3A’s domains is critical to dissecting its role in disease and identifying novel therapeutic strategies. Although x-ray crystallography and cryo-electron microscopy have yielded deep mechanistic insight into DNMT3A regulation^14,17,21–23^, to date no published structure has successfully resolved all three domains. A recent study evaluated the functional consequences of clinically observed mutations across DNMT3A’s full sequence^24^, demonstrating the potential for large-scale mutational analysis to reveal novel insight. While this study used exogenously expressed DNMT3A, approaches such as CRISPR scanning could provide the opportunity to conduct high-throughput mutational screening directly within the endogenous gene^25–28^. This approach has illuminated functional mechanisms of other chromatin regulators^25,29^, but is less easily applied to proteins like DNMT3A whose loss is tolerated in cultured cells^30^. This is because Cas9 predominately generates frameshift mutations that can obscure in-frame mutations of interest, unless actively removed from the gene pool due to a fitness disadvantage. Moreover, the resulting in-frame mutations can be highly complex, necessitating time-consuming clonal analysis for their functional dissection^25,29^. In this regard, cytosine base editors, which avoid insertion-deletions (indels) and have more finely targeted and predictable mutational outcomes^31^, greatly empower CRISPR mutational scanning. This technology, however, has so far been limited to viability screens^32–35^.

We envisioned that base editors could enable high-throughput interrogation of DNMT3A in its native state. By applying a genetically encoded methylation reporter, we expand the scope of base editor screens and map functional hotspots throughout DNMT3A encompassing disease-associated residues. We use biochemical analysis to characterize an ADD-MTase interface that is critical for allosteric activation. We also identify mutations in the PWWP domain that affect DNMT3A function, yet surprisingly not through destabilization or abrogation of histone targeting. Instead, these mutations impact DNA binding by the PWWP domain, which we show through biochemical and genomic approaches is a noncanonical function required for full DNMT3A activity. More broadly, our study demonstrates a generalizable approach to studying essential and nonessential chromatin regulators.

## Results

### A live-cell reporter enables fluorescence-based readout of endogenous DNMT3A activity

To enable readout of endogenous DNMT3A activity, we leveraged a chromatin regulation reporter that provides a fluorescence-based readout of gene silencing activity^36^. We fused the accessory factor DNMT3L, which itself is catalytically inactive^12^, to the reverse Tet repressor (rTetR) in order to bind endogenous DNMT3A and recruit it to a genome-integrated reporter cassette in a doxycycline (dox)-inducible fashion (**Fig. 1a and Extended Data Fig. 1a**). Subsequent DNA methylation of the reporter promoter would silence citrine expression, thus providing a fluorescence-based readout of DNMT3A activity amendable to base editor scanning (**Fig. 1b**). We generated a clonal K562 reporter cell line and validated its ability to report on DNA methylation silencing. Upon treating these cells with dox, we observed complete reporter silencing with clearly resolved silenced and unsilenced states (**Fig. 1c,d and Extended Data Fig. 1b**). Silenced cells did not revert after dox washout, consistent with the stability of gene repression due to DNA methylation^36^ (**Fig. 1c**). Moreover, targeted bisulfite sequencing of the integrated reporter showed a time-dependent gain of promoter methylation in dox-treated cells, confirming that methylation accompanies silencing (**Extended Data Fig. 1c**).

**Fig. 1.**
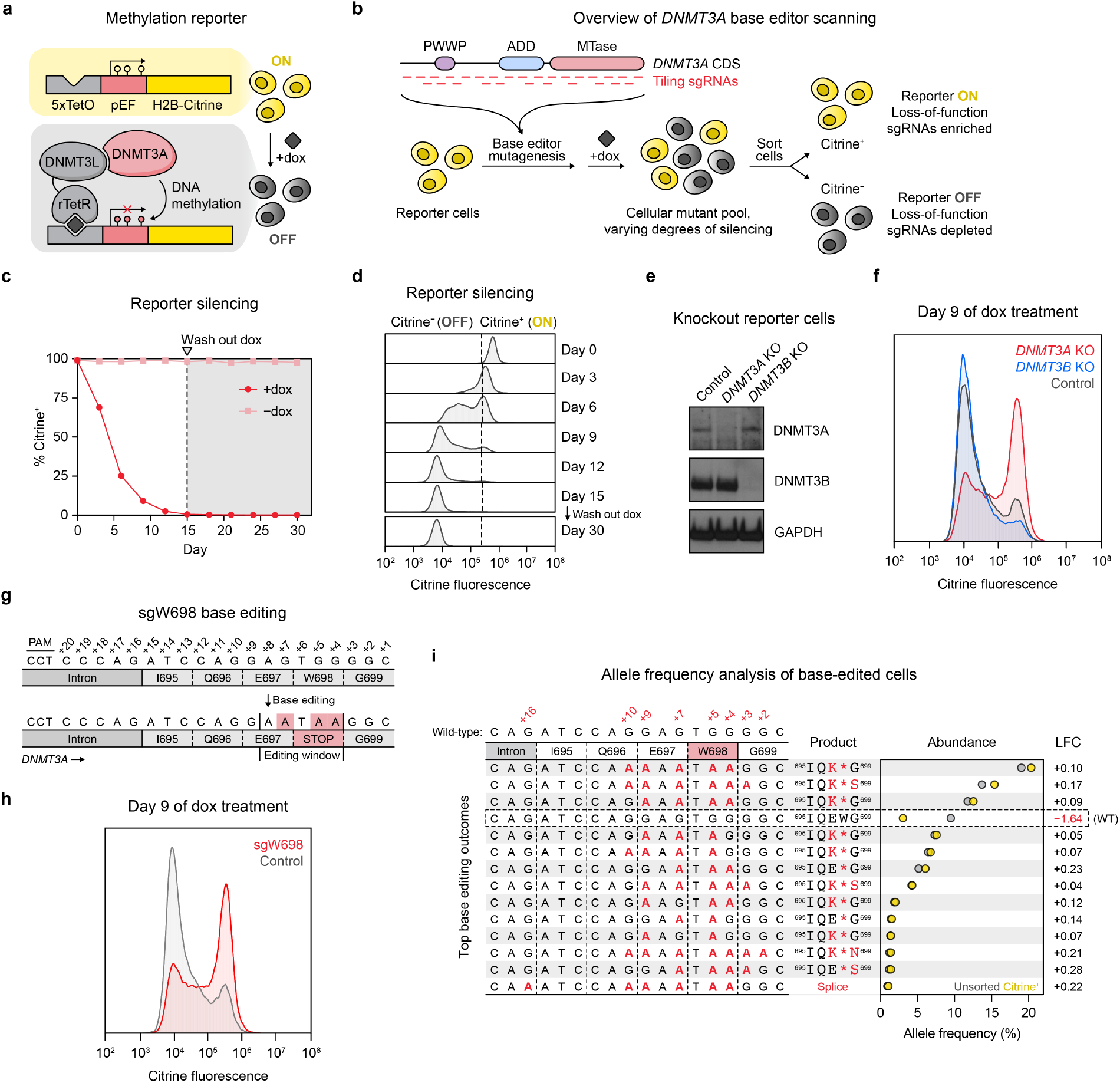
A live-cell reporter enables fluorescence-based readout of endogenous DNMT3A activity. **a**. Schematic of the methylation reporter. rTetR, reverse tetracycline repressor; TetO, tetracycline operator; pEF, *EF1*α promoter. Filled lollipops indicate DNA methylation. **b**. Overview of *DNMT3A* base editor scanning experiment. **c**. Timecourse of reporter silencing measured by flow cytometry. Dotted line indicates dox washout on day 15. Data are mean ± SD of n = 3 replicates. **d**. Citrine fluorescence of dox-treated cells at indicated timepoints from **c.** **e**. Immunoblot of *DNMT3*-knockout reporter cells. Control, sgRNA recognizing the luciferase coding sequence (nontargeting *LucA* sgRNA). Uncropped images are provided in Supplementary Fig. 1. **f**. Citrine fluorescence of cells from **e** measured by flow cytometry after 9 d of dox treatment. **g**. sgW698 target site in *DNMT3A*. Numbers indicate positions along the protospacer (antisense to the *DNMT3A* gene). Expected base editing mutations are highlighted in red (these appear as G to A because the protospacer is along the opposite strand). **h**. Citrine fluorescence of cells treated with sgW698 or control measured by flow cytometry after 9 d of dox treatment. **i**. Allele frequencies in cells edited with sgW698 after 9 d of dox treatment, either sorted for citrine^+^ cells (yellow dots) or unsorted (gray dots). The wild-type allele is boxed, and only alleles with ≥1% allele frequency in at least one sample are shown. Protein product sequences are shown with nonsynonymous mutations in red. Splice, splice site mutation; LFC, log_2_(fold-change allele frequency in citrine^+^ versus unsorted cells). Genotyping was performed once. Histograms of citrine fluorescence in **d, f**, and **h** are representative of n = 3 replicates. One of two independent experiments is shown for experiments in **c–f** and **h**.

We next evaluated the sensitivity of this reporter toward DNMT3A loss-of-function mutations, which should impair its silencing under dox treatment. Because mammalian cells possess a second de novo DNMT, DNMT3B, we first tested whether this might interfere with our ability to detect DNMT3A-mediated silencing. We found that CRISPR-Cas9 knockout of *DNMT3A*, but not *DNMT3B*, caused defective reporter silencing (**Fig. 1e,f and Extended Data Fig. 1d**), as expected since K562 cells predominantly express inactive isoforms of DNMT3B (ref. ^37^). This showed that our reporter selectively detects loss of DNMT3A. To assess sensitivity toward inactivating C to T base editing mutations, we designed a single-guide RNA (sgRNA) with the goal of introducing a nonsense mutation at the W698 codon, named sgW698 (**Fig. 1g**). Reporter cells edited with sgW698 displayed a silencing defect similar to that of *DNMT3A*-knockout cells (**Fig. 1h and Extended Data Fig. 1e**). We isolated cells remaining citrine^+^ using fluorescence-activated cell sorting (FACS) and deep sequenced the edited site, finding high rates of base editing and low rates of indels (**Extended Data Fig. 1f**). Relative to unsorted cells, citrine^+^ cells were markedly depleted of the unedited, wild-type allele and enriched with alleles containing the W698* mutation (**Fig. 1i**), showing that inactivating *DNMT3A* mutations are indeed enriched in cells defective for silencing. Taken together, these results demonstrate that our reporter can detect base editing mutations introduced in the endogenous *DNMT3A* gene, enabling mutational scanning of the native protein.

### Base editor scanning charts mutations across DNMT3A that impact function

To carry out base editor scanning, we designed a library containing all sgRNAs targeting exons and flanking intronic sequences of DNMT3A isoform 2 (DNMT3A2), the predominant isoform expressed in K562 (ref. ^37^). We transduced this library into reporter cells, treated the cells with dox, and sorted them based on citrine fluorescence (**Fig. 1b**). We used deep sequencing to quantify sgRNA abundances in citrine^+^ and citrine^−^ fractions, comparing these to unsorted cells to define a metric for whether a particular sgRNA was enriched or depleted upon FACS sorting (termed “sgRNA score,” see **Methods**). Analysis of these sgRNA scores showed that sgRNA abundances were inversely correlated between citrine^+^ and citrine^−^ fractions, as expected (**Extended Data Fig. 2a,b**). In particular, sgRNAs that were enriched in citrine^+^ cells at day 9 (cells that were resistant to silencing) were conversely depleted in citrine^−^ cells at day 3 (cells that began to silence early) (**Extended Data Fig. 2b**). This dichotomy and the broad correlations we observed provided confidence in our base editor scanning experiment.

To enable further analysis, we classified each sgRNA based on its predicted mutational outcome, assuming complete editing within the canonical +4 to +8 editing window^31^. Focusing on citrine^+^ cells at day 9, we found that sgRNAs predicted to introduce nonsense or splice site-disrupting mutations (referred to as “nonsense sgRNAs” or “splice site sgRNAs,” respectively) were largely enriched (positive sgRNA score), consistent with loss-of-function (**Fig. 2a,b**). Conversely, most silent and intronic sgRNAs were unchanged in abundance, as expected. However, missense sgRNAs exhibited a wide range of scores, reflecting the context-dependent effects of missense mutations. For example, those targeting residues at the active site or near the DNMT3A-DNMT3L binding interface had higher sgRNA scores on average compared to other missense sgRNAs (**Extended Data Fig. 2c**). In general, most hit missense sgRNAs (ones that were enriched or depleted), mapped to within protein domains (**Fig. 2a and Extended Data Fig. 2d**).

**Fig. 2.**
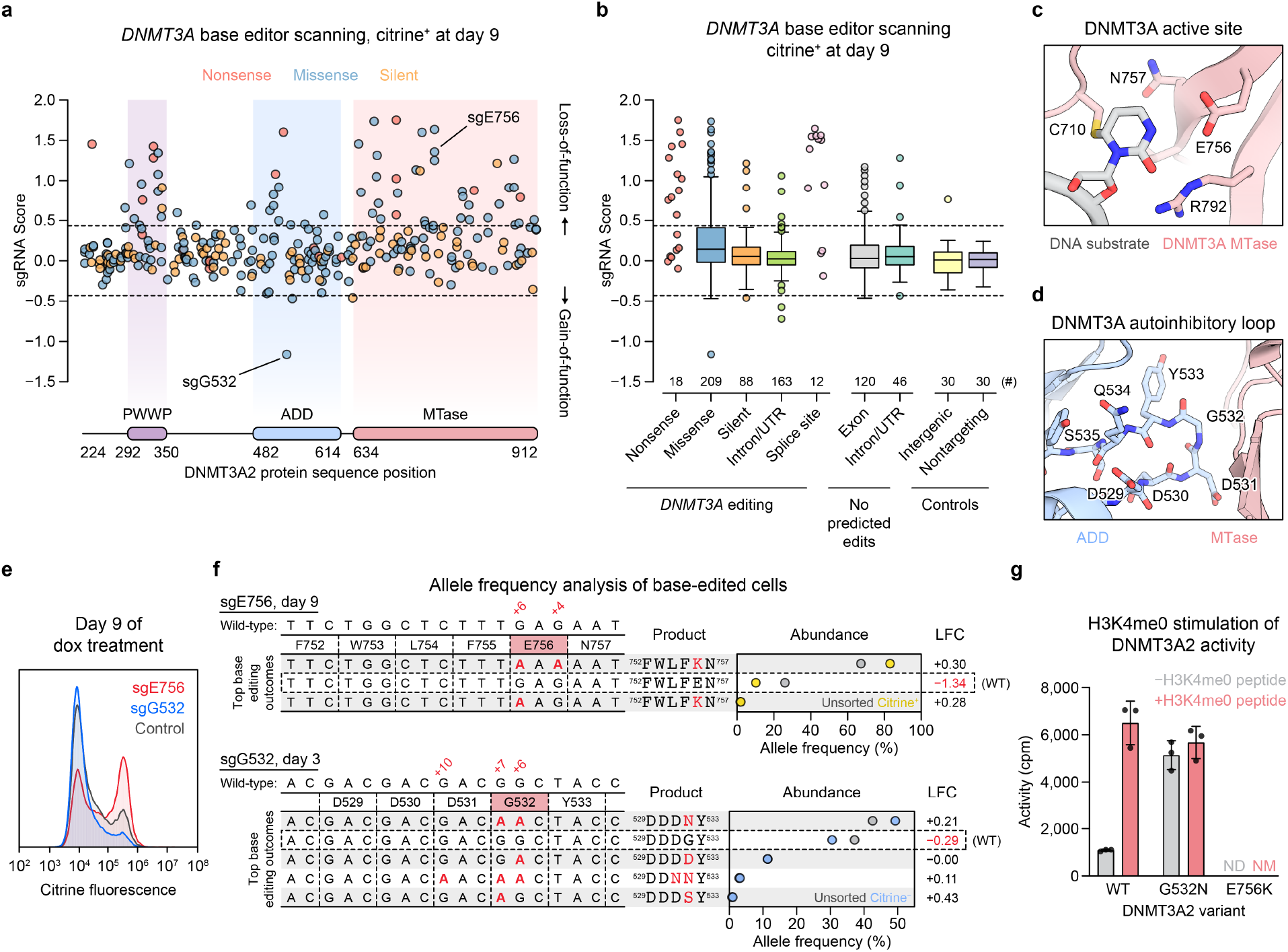
Base editor scanning charts mutations across DNMT3A that impact function. (**a–b**) *DNMT3A* base editor scanning results for citrine^+^ cells at day 9. Dotted lines indicate ±2 SD of intergenic control sgRNAs, and data are the average of n = 3 replicates. **a**. Scatterplot of sgRNA scores for nonsense (red), missense (blue), and silent (orange) sgRNAs, plotted against the targeted site in the coding sequence. **b**. Boxplot of sgRNA scores for sgRNAs classified by predicted editing outcome. Center line, median; box, interquartile range; whiskers, up to 1.5 × interquartile range per the Tukey method. Outliers and any categories with n < 20 are shown individually. The number of sgRNAs in each category is printed below the box. **c**. View of the DNMT3A active site (red) with the DNA substrate (gray) (PDB: 5YX2). **d**. View of the ADD (blue)-MTase (red) autoinhibitory interface highlighting the autoinhibitory loop (PDB: 4U7P). **e**. Citrine fluorescence of base editor-treated cells measured by flow cytometry after 9 d of dox treatment. Control, nontargeting *LucA* sgRNA. Histograms are representative of n = 3 replicates. **f**. Allele frequencies in base-edited cells. Top, cells edited with sgE756 after 9 d of dox treatment, comparing citrine^+^ (yellow dots) to unsorted (gray dots) cells. Bottom, cells edited with sgG532 after 3 d of dox treatment, comparing citrine^−^ cells (blue dots) to unsorted cells (gray dots). The wild-type allele is boxed, and only alleles with ≥1% allele frequency in at least one sample are shown. Protein product sequences are shown with nonsynonymous mutations in red. LFC, log_2_(fold-change allele frequency in sorted versus unsorted cells). Genotyping was performed once. **g**. Activity of purified DNMT3A2 in the presence (red) or absence (gray) of H3K4me0 peptide. Data are mean ± SD of n = 3 replicates. ND, not detected; NM, not measured. One of two independent experiments is shown for experiments in **e** and **g**.

To validate the results of our screen, we studied two sgRNAs with opposing scores, sgE756 and sgG532 (**Fig. 2a**). sgE756 was one of several highly enriched sgRNAs predicted to mutate E756, a key residue in DNMT3A’s catalytic mechanism^38^ (**Fig. 2c**), thereby causing loss-of-function. By contrast, sgG532 was strongly depleted in citrine^+^ cells, consistent with gain-of-function. Because G532 is in the ADD domain autoinhibitory loop (**Fig. 2d**), we hypothesized that sgG532 might activate DNMT3A by disrupting autoinhibition. Testing each sgRNA individually, we found that sgE756-treated reporter cells were defective in silencing relative to control, while sgG532-treated cells displayed greater silencing (**Fig. 2e and Extended Data Fig. 2e**). We genotyped sgE756-treated cells remaining citrine^+^ at day 9, finding that the sorted cells were depleted of wild-type alleles in favor of alleles with the E756K mutation (**Fig. 2f and Extended Data Fig. 2f**). For sgG532, we considered cells that had already begun to silence at day 3 since these should be enriched for gain-of-function mutations; indeed, citrine^−^ sgG532-treated cells were enriched for the G532N allele. To confirm that the E756K and G532N mutations indeed alter catalytic activity, we purified recombinant DNMT3A2 (**Extended Data Fig. 3a**) and conducted enzyme activity assays. Indeed, the E756K mutant was catalytically dead, while the G532N mutant showed increased activity relative to wild-type (**Fig. 2g**). H3K4me0 peptide is known to promote the release of DNMT3A autoinhibition^17^. Unlike wild-type DNMT3A2, we found that DNMT3A2 G532N could not be further stimulated by the addition of H3K4me0 peptide, suggesting that its hyperactivity arises due to blocking the autoinhibited conformation. Taken together, these results demonstrate that our base editor scanning approach can accurately discern mutations that perturb DNMT3A function.

### Functional hotspot analysis highlights an interdomain interface important for allosteric activation

Base editor scanning identified enriched missense sgRNAs across the *DNMT3A* coding sequence. Since functional regions in proteins may comprise residues that are far apart in the linear sequence, we considered whether analyzing these sgRNAs, and their targeted residues, in three-dimensional (3D) space might reveal shared mechanisms. We focused on missense sgRNAs targeting the ADD and MTase domains, since structures have been solved for this portion in both active and autoinhibitory conformations^17^. We began by calculating a proximity-weighted enrichment score (PWES)^25^ for each pair of sgRNAs describing their 3D proximity and combined sgRNA scores (see **Methods**). To do so, we mapped each sgRNA to its target residue in the active conformation, reasoning that this would enable us to identify functionally critical areas whose mutation causes loss-of-function. Hierarchical clustering of the resulting PWES matrix defined eight clusters of sgRNAs (**Fig. 3a**). The two most highly enriched clusters (**Fig. 3b**) corresponded to known functional hotspots: cluster 3 targeted the active site, while cluster 4 spanned the active site and RD homodimer interface, which is important for catalytic activity^39^ (**Fig. 3c**). We validated sgRNAs from these clusters (sgD641, sgE664/V665, and sgD668) by individual transduction into reporter cells, verifying on-target C to T editing, minimal indels, and the expected reporter silencing defect (**Extended Data Fig. 4a–d**). Clusters 3 and 4 together contained most of the highest-scoring ADD and MTase domain missense sgRNAs, with the notable exception of sgL737, a cluster 1 sgRNA that we found mutated an α-helix at the DNMT3A-DNMT3L interface—thus presumably disrupting recruitment of DNMT3A to the reporter locus. Altogether, these results support the utility of base editor scanning with PWES analysis to identify critical functional hotspots.

**Fig. 3.**
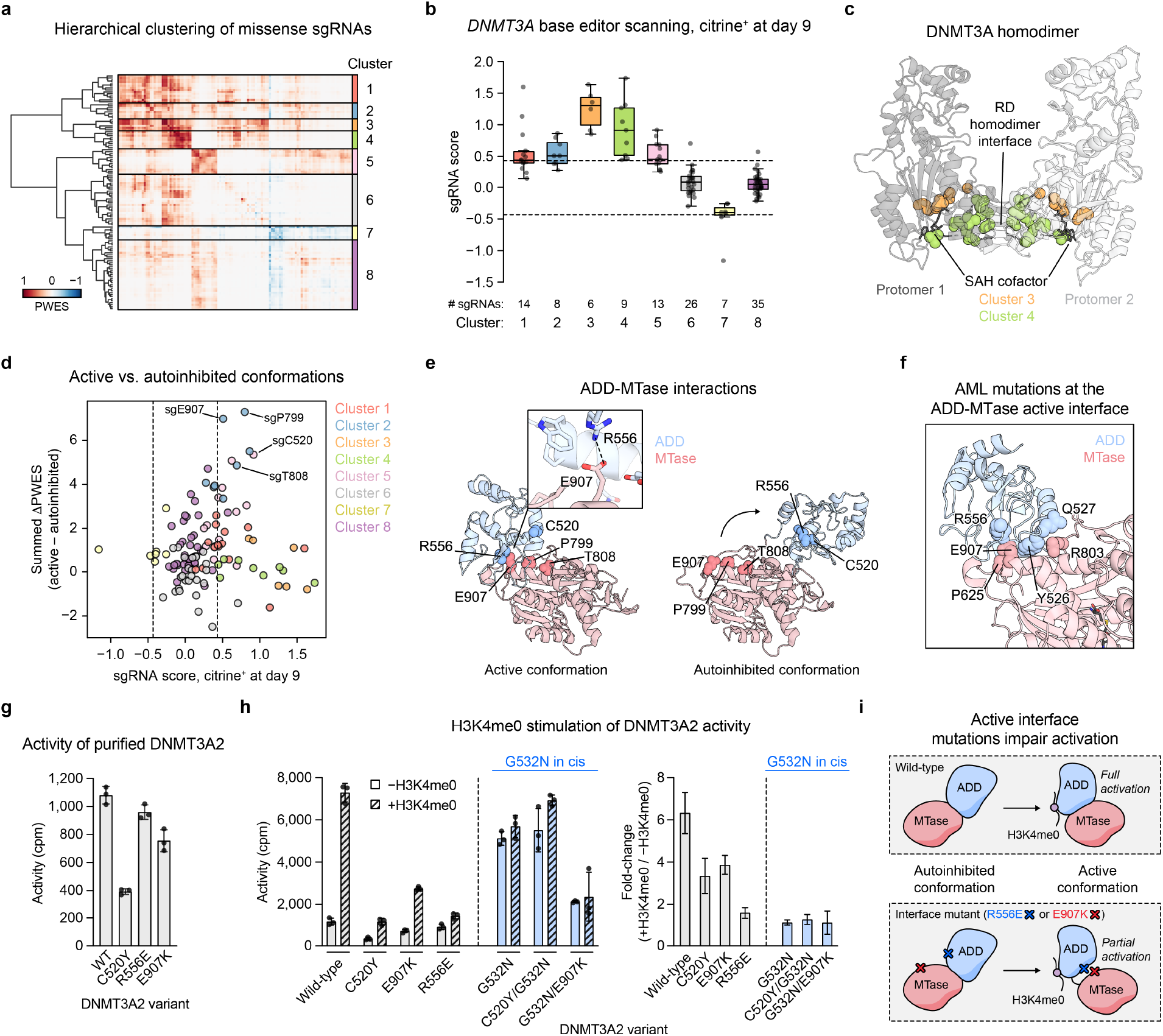
Functional hotspot analysis highlights an interdomain interface important for allosteric activation. **a**. Heatmap depicting the PWES matrix for all pairs of missense sgRNAs (n = 118) mapping to resolved residues in the structure of active conformation DNMT3A (PDB: 4U7T). sgRNAs are ordered by hierarchical clustering. **b**. Boxplot showing sgRNA scores in citrine^+^ cells at day 9 of the base editor scanning experiment, with sgRNAs organized by clusters from **a**. Data are the average of n = 3 replicates. Dotted lines indicate ±2 SD of intergenic control sgRNAs. Center line, median; box, interquartile range; whiskers, up to 1.5 × interquartile range per the Tukey method. Individual data points are overlaid. **c**. View of the DNMT3A homodimer with residues targeted by sgRNAs in clusters 3 and 4 highlighted (PDB: 4U7T). **d**. Comparison of PWES calculated using active conformation (PDB: 4U7T) 3D proximity versus that using autoinhibited conformation (PDB: 4U7P) 3D proximity, represented as the summed ΔPWES (see Methods for details). sgRNA scores, colors, and dotted lines correspond to those in **b.** **e**. View of the ADD (blue)-MTase (red) interface in the structures of DNMT3A in active (left, PDB: 4U7T) or autoinhibited (right, PDB: 4U7P) conformations. Inset shows the ionic bond mediated by R556 and E907. **f**. View of active conformation DNMT3A highlighting residues at the ADD-MTase interface that are mutated in AML (data from COSMIC) (PDB: 4U7T). **g**. Activity of purified DNMT3A2. Data are mean ± SD of n = 3 replicates. **h**. Stimulation of purified DNMT3A2 by H3K4me0 peptide. Right, fold-change in activity in the presence of H3K4me0 versus in the absence of H3K4me0. Data are mean ± SD of n = 3 replicates. Fold-change errors were propagated from the individual SDs. One of two independent experiments is shown. **i**. Cartoon depicting impaired H3K4me0 stimulation caused by active conformation ADD-MTase interface-disrupting mutations.

We next assessed whether our results could also nominate novel regions of regulatory importance. We hypothesized that computing the PWES values using the structure of autoinhibited DNMT3A and then comparing them to those obtained with the active conformation might highlight residues playing key allosteric roles. To test this, we calculated a “summed ΔPWES” for each sgRNA representing this difference (see **Methods**). For most sgRNAs, this measure was close to zero, indicating that their proximity-weighted correlation to other sgRNAs is not greatly altered by the conformational change (**Fig. 3d**). However, several sgRNAs in cluster 2 (sgP799, sgT808, and sgE907) and cluster 5 (sgC520) displayed large, positive summed ΔPWES values, suggesting that the transition from the autoinhibitory to the active conformation strengthens their correlations with other sgRNAs. These sgRNAs mapped to residues at or near the active ADD-MTase interface (**Fig. 3e**), suggesting that it might play a functionally important role. Indeed, mutations at this interface have been identified in AML patients^40^ (**Fig. 3f**), suggesting this may be a clinically relevant functional hotspot. We validated these sgRNAs in reporter cells, finding on-target editing but subtle effects (sgP799, sgT808, and sgE907) on reporter silencing (**Extended Data Fig. 4a– d**). However, given the incomplete silencing defect we observed even for catalytically dead DNMT3A E756K (**Fig. 2e**), we reasoned that the rapid timescale of reporter silencing, as well as the presence of heterozygous cells harboring wild-type DNMT3A, likely hindered our ability to detect partial loss-of-function phenotypes.

To more directly test the importance of the active ADD-MTase interface, we considered the impact of mutations at this interface on catalytic activity. Since E907 stabilizes this interface through an ionic bond with R556 (**Fig. 3e**), we tested the E907K (product of sgE907) and R556E charge-reversal mutants. Surprisingly, neither the E907K nor the R556E mutant had greatly reduced activity (**Fig. 3g**). Both, however, impaired the release of autoinhibition by H3K4me0, as measured by a lower fold-change stimulation than wild-type upon addition of peptide (**Fig. 3h**). As a comparator, we tested the C520Y mutant (predicted product of sgC520) since C520 is not directly at this interface. Although genotyping of base-edited cells revealed strong editing of a splice site outside the editing window (+9 position) (**Extended Data Fig. 4b**), we nonetheless found that the DNMT3A2 C520Y was catalytically defective and displayed impaired stimulation (**Fig. 3g**).

We next considered whether blocking the autoinhibitory conformation by the G532N mutation in cis, as an orthogonal mode of activation, could instead rescue full activity. The C520Y/G532N double mutant was comparably active to the G532N single mutant, even though the C520Y single mutant was not fully stimulated by H3K4me0 (**Fig. 3h**). This suggests that although the C520Y mutation interferes with allosteric activation, its effects can be overcome. By contrast, the G532N/E907K double mutant displayed less activity than the G532N single mutant, consistent with the incomplete H3K4me0 stimulation observed for the E907K single mutant. Thus, E907K causes loss-of-function through a distinct mechanism than C520Y. Taken together, these results suggest that the active ADD-MTase interface, which is stabilized by the E907-R556 interaction, is necessary for full activation. Promoting the autoinhibitory to active conformational change— either by H3K4me0 binding or by the G532N mutation in cis—is not sufficient for maximal activity in the absence of additional interactions specific to the active conformation (**Fig. 3i**). While mutations preserving H3K4me0 binding but abrogating stimulation have been reported^41^, these effects have not previously been tied directly to the ADD-MTase interface. Moreover, both E907 and R556 are mutated in AML^40^, suggesting that this loss-of-function pathway is clinically relevant. Our mechanistic analysis thus finds that the active ADD-MTase interface is a functional hotspot, demonstrating the power of leveraging base editor scanning with 3D structural analysis to uncover mechanistic insight.

### PWWP domain mutations have variable impacts on protein stability

We next turned our attention to the sgRNA hits within the PWWP domain (**Fig. 4a,b**). Because our methylation reporter involves artificial DNMT3A recruitment, loss of the canonical H3K36me2 reader function of the PWWP domain^18,19^ could not by itself explain enrichment in our screen. We thus set out to survey these hits, beginning by validating a panel of sgRNAs by individual transduction into reporter cells. Genotyping confirmed the presence of C to T editing around the targeted sites in *DNMT3A*, concomitant with dramatic defects in reporter silencing for sgG293/E294, sgS337.1, and sgS337.2, as well as smaller but detectable effects for sgR301 and sgS312.2 (**Figs. 4c,d**). Among these hits were also two sgRNAs annotated as causing no predicted edits (sgS312.1) or only a silent mutation (sgE342.2) (**Fig. 4a**), whose effects we hypothesized might arise from editing outside of the predicted +4 to +8 window. Subsequent genotyping confirmed that these sgRNAs produce the corresponding mutations, S312F and E342K, by editing at the +9 and +10 positions, respectively (**Fig. 4c**). We note that sgE342.1 and sgR366 did not positively validate in one of two independent trials.

**Fig. 4.**
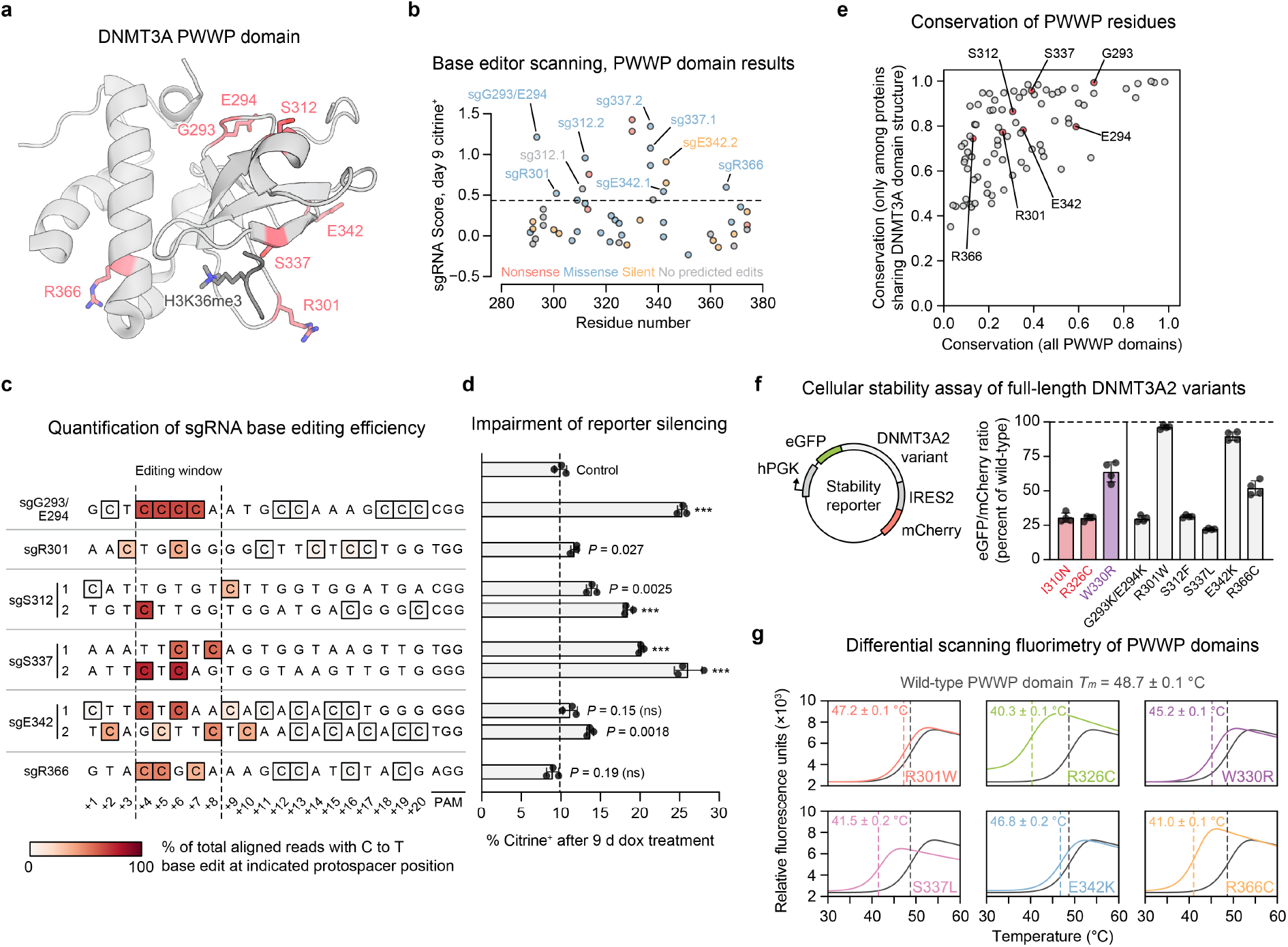
PWWP domain mutations have variable impacts on protein stability. **a**. View of the DNMT3A PWWP domain (gray) with key residues targeted by enriched missense sgRNAs shown as red sticks (PDB: 3LLR; H3K36me3 (dark gray) aligned from 5CIU). **b**. Summary of base editor scanning results within the PWWP domain. Dotted line indicates mean + 2 SD of intergenic control sgRNAs. Data are the same as in Fig. 2a and are the average of n = 3 replicates. **c**. Base editing efficiency for selected PWWP sgRNA hits. Each row depicts a protospacer sequence. The heatmap intensity shows the efficiency of C to T editing at each C nucleotide, as measured by deep sequencing. Genotyping was performed once. **d**. Flow cytometric quantification of cells edited with PWWP sgRNAs or *LucA* nontargeting control remaining unsilenced after 9 d of dox treatment. Data are mean ± SD for n = 3 replicates. *P* values (***, *P* ≤ 0.001; ns, not significant) were calculated through two-tailed unpaired *t* tests comparing each sgRNA to control. **e**. Conservation of DNMT3A PWWP domain residues among all PWWP domains (horizontal axis) and among only proteins sharing the domain structure of DNMT3A (vertical axis). **f**. Stability of DNMT3A2 variants measured by an eGFP-mCherry fluorescence reporter in K562 cells (schematic shown to left). Data are mean ± SD of n = 4 measurements (two measurements of each of two independently transduced cell lines). Colored bars represent previously reported unstable (red) or stable (purple) disease-associated mutants^5,24^. **g**. Thermal stability of purified PWWP domain variants measured by differential scanning fluorimetry. The wild-type curve (gray) is superimposed over that of each mutant. Curves represent mean of n = 3 replicates. Melting temperatures (*T*_*m*_) are printed for each variant (mean ± SD) and indicated by colored dashed lines. For **d** and **g**, one of two independent experiments is shown.

Having confirmed DNMT3A loss-of-function with several sgRNAs, we considered whether these effects might be due to disrupting structural integrity, as opposed to disrupting allosteric processes. Indeed, recent work has found that a large fraction of PWWP clinical mutations cause destabilization and degradation^5,24^, suggesting this is a common loss-of-function pathway. To narrow our list of hits, we investigated the evolutionary conservation of their targeted residues (**Fig. 4e**). We reasoned that residues universally conserved across PWWP domain-containing proteins likely serve important structural roles, and thus might be structurally intolerant to mutation. By contrast, residues conserved only in DNMT3A-like proteins might play functional roles specific to DNMT3A. Both G293 and E294 were highly conserved across proteins with PWWP domains, suggesting that sgG293/E294 might cause destabilization. However, the majority of validated sgRNAs targeted residues that were not conserved as broadly, namely R301, S312, S337, and E342.

To experimentally test whether these mutations affect protein stability in cells, we studied DNMT3A2 expression in K562 cells using a dual-fluorescence stability reporter^42^ (**Fig. 4f**). Here, eGFP is fused to DNMT3A2 so that its fluorescence is a proxy for the amount of DNMT3A2 in the cell. mCherry is expressed co-transcriptionally so that the ratio of eGFP to mCherry fluorescence provides a normalized measure of DNMT3A2 levels (**Extended Data Fig. 5**). To validate this assay, we tested several disease-associated mutations whose effects on stability have been characterized. Consistent with previous reports^5,24^, both the I310N TBRS and R326C clonal hematopoiesis mutations were destabilizing, reducing DNMT3A2 levels to about 30% of wild-type levels, while the stable W330R HESJAS mutation preserved over 60% of wild-type expression (**Fig. 4f**). As expected based on our conservation analysis, the G293K/E294K mutant was similarly unstable as the I310N mutant. However, we observed variable stability for the remaining mutants. The S312F and S337L mutants were unstable, which in light of our conservation analysis suggest that S312 and S337 play structural roles specific to the DNMT3A architecture. Conversely, the R301W and E342K mutants were nearly as stable as wild-type, and in fact were expressed at higher levels than the W330R mutant.

Although cellular expression is a useful proxy for protein stability, we next sought to rigorously evaluate the impacts of these mutations on biochemical stability. To do so, we purified recombinant PWWP domains (**Extended Data Fig. 3b**) and performed differential scanning fluorimetry to assay for thermal stability (**Fig. 4g**). These results were highly concordant with the cellular stability reporter data. In particular, the R326C, S337L, and R366C mutants all had a greatly lowered melting temperature, indicating that they are intrinsically less stable than wild-type. Interestingly, the instability of the R366C mutant was more pronounced here than in our stability reporter assay (**Fig. 4f**), suggesting that its biochemical instability may be attenuated in the context of full-length DNMT3A2 in cells. Incomplete destabilization in cells could explain why sgR366 failed to validate in our reporter silencing assay (**Fig. 4d**), as lowered levels of DNMT3A2 might still support full reporter silencing. By contrast, the melting temperatures of the R301W and E342K mutants were similar to that of wild-type (**Fig. 4g**). This confirmed that these mutations, both of which have been identified in disease^24,43^, do not adversely impact protein stability. Taken together, these results show that DNMT3A loss-of-function indeed arises from destabilizing PWWP domain mutations, yet also point to the existence of additional mechanisms by which the stable R301W and E342K mutations impact function.

### DNA binding by the PWWP domain modulates DNMT3A activity

Given that neither the R301W nor E342K mutations impact DNMT3A stability, we investigated whether they might cause loss-of-function by directly impairing catalysis. Activity assays showed opposing effects for these mutants, with the R301W mutant losing catalytic activity, and the E342K mutant surprisingly gaining activity (**Fig. 5a**). We considered whether these mutations might impact the PWWP domain’s canonical histone reader function^18,19^, and the recent reports of stimulation by H3K36me2 (ref. ^19,20^). Testing for stimulation by the H3 peptide (residues 21– 44), we found that wild-type DNMT3A2 was stimulated selectively by H3K36me2 but not by H3K36me0. As a control, we tested the W330R mutant, which lacks histone reader function due to a binding pocket mutation (ref. ^5^). DNMT3A2 W330R was refractory to histone stimulation, and moreover, was hyperactive at baseline like DNMT3A2 E342K. With regards to the R301W mutant, we observed stimulation by H3K36me2 selectively over H3K36me0 and to a similar degree as wild-type, suggesting it retains H3K36me2 recognition. On the other hand, the activity of E342K variant was not greatly altered by addition of either peptide, though its basal activity was comparable to that of stimulated wild-type enzyme; in other words, E342K appeared to pre-activate DNMT3A.

**Fig. 5.**
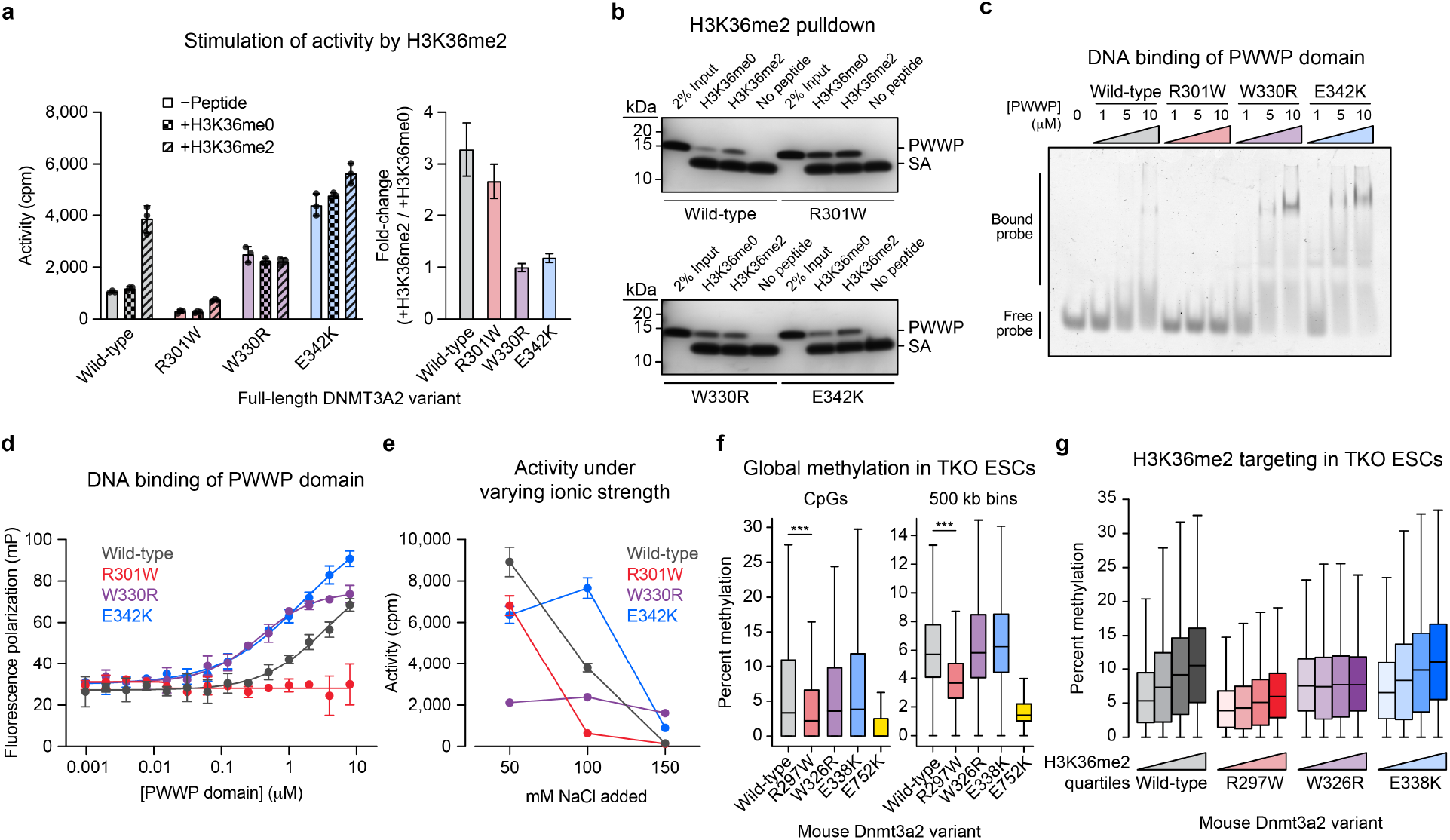
DNA binding by the PWWP domain modulates DNMT3A activity. **a**. Stimulation of DNMT3A2 by H3K36me2 peptide (residues 21–44). Left, activity of purified DNMT3A2 variants in the presence of H3K36me0 or H3K36me2 peptides, or in the absence of either peptide. Right, fold-change in activity observed with H3K36me2 compared to with H3K36me0. **b**. Binding of purified PWWP domain variants to biotinylated H3K36me0 or H3K36me2 peptides. Bound proteins were captured by streptavidin pulldown, resolved by SDS-PAGE, and visualized by silver staining. SA, streptavidin. **c**. Binding of purified PWWP domains to a Cy3-labeled 30 bp oligonucleotide probe measured by electrophoretic mobility shift assay. Image was inverted and brightness was adjusted for visualization. **d**. Binding of purified PWWP domains to a Cy3-labeled 30 bp oligonucleotide probe measured by fluorescence polarization assay. **e**. Activity of purified DNMT3A2 variants under increasing ionic strength. Reactions were conducted in a low ionic strength assay buffer supplemented with the indicated concentrations of NaCl (see Methods for additional details). Data in **a, d**, and **e** are mean ± SD for n = 3 replicates. Fold-change errors in **a** were propagated from the individual SDs. For **a–e**, one of two independent experiments is shown in each panel. Unprocessed images for data shown in **b** and **c** are provided in Supplemental Fig. 1. **f**. CpG methylation genome-wide in TKO ESCs ectopically expressing the indicated Dnmt3a2 variants. Left, CpG methylation for all CpGs, excluding CpGs with zero methylation in all samples (n = 683,371). Right, CpG methylation averaged across 500 kb bins (n = 5,204). *P* values (***, *P* < 2.3 × 10^−308^) were calculated through two-sided Wilcoxon signed-rank tests. **g**. CpG methylation within 10 kb genomic bins ranked into quartiles based on normalized H3K36me2 ChIP-seq signal (n = 93,516 bins total, 23,379 bins per quartile). For **f–g**, only CpGs with 5× coverage across all samples were considered. Methylation values represent the average of two biological replicates.

The hyperactivity of the E342K variant in biochemical assays was unexpected, since sgE342.2 registered as loss-of-function in our reporter assay (**Fig. 4d**). Since sgE342.2 edited outside of the editing window (**Fig. 4c**), we considered whether bystander mutations might account for this discrepancy. We performed genotyping after sorting for citrine^+^ cells, which enriches for loss-of-function mutations, testing several sgRNAs targeting the E342 codon (**Extended Data Fig. 6a**). Counterintuitively, the E342K allele was not correlated with impaired function, as it was depleted in the citrine^+^ fraction for both sgE342.1- and sgE342.2-treated cells and abundant in cells treated with sgE342.3, a third sgRNA that was not enriched in our screen (**Extended Data Fig. 6b**). For sgE342.2, sorting for citrine^+^ cells instead enriched for alleles containing E342K and additional silent mutations, suggesting that reporter loss-of-function may have arisen from synonymous mutations impairing expression. Nonetheless, we decided to further investigate both the E342K and R301W mutants biochemically due to their stark differences.

To more directly assess whether the E342K mutation impacts discrimination between H3K36 methylation states, we conducted a pulldown assay measuring binding of purified PWWP domains to a biotinylated H3 peptide. Wild-type PWWP domain selectively bound H3K36me2 over H3K36me0, while PWWP W330R showed no preference, consistent with previous studies^5,44^ (**Fig. 5b**). We observed high signal for the PWWP R301W with both histone modification states, possibly suggesting loss of binding specificity. However, we reasoned that by further increasing the net negative charge, the R301W mutation might promote nonspecific binding to the positively charged histone peptide. Intriguingly, PWWP E342K bound H3K36me2 stronger than H3K36me0, indicating that it retains H3K36me2 recognition, despite DNMT3A2 E342K’s lack of stimulation by H3K36me2 (**Fig. 5a**). These results suggest that the R301W and E342K mutations affect function in opposing ways, and through a mechanism distinct from H3K36me2 recognition.

Considering the crystal structure of the DNMT3A PWWP domain, we noticed that both the R301W and E342K mutations alter charged surface residues (**Fig. 4a**). Thus, we hypothesized that their opposing effects on enzyme activity might be due to increased or decreased binding of negatively charged DNA. Indeed, prior studies have demonstrated that the PWWP domain nonspecifically binds DNA^44,45^, though this phenomenon has not been shown to contribute positively to DNMT3A function. Thus, we conducted electrophoretic mobility shift and fluorescence polarization assays to test binding of the purified PWWP domain to a DNA oligonucleotide probe (**Fig. 5c,d**). Both assays confirmed a loss of DNA affinity for the R301W mutant, and gain for the E342K variant, consistent with expectations based on the net changes in charge. We also observed increased DNA affinity for the W330R variant, similar to the E342K variant. This was consistent with its increased net positive charge, and moreover suggested a potential link between DNA binding and the increased basal activity of the W330R and E342K mutants (**Fig. 5a**).

To test whether altered DNA binding indeed accounts for these differences in catalytic activity, we measured the mutants’ activity profiles under varying ionic strength. A similar assay was used to characterize a DNA binding site in DNMT3A’s N-terminus^46^, since the charged interactions underlying DNA affinity are attenuated by increased ionic strength. We supplemented a low ionic strength assay buffer with increasing levels of NaCl (see **Methods**). Compared to wild-type, the R301W mutation sensitized DNMT3A2 to increasing ionic strength, while both the W330R and E342K mutations conferred insensitivity, consistent with weaker and stronger DNA binding, respectively (**Fig. 5e**). Interestingly, the hyperactive nature of the W330R and E342K mutants was only apparent at higher ionic strengths, similar to our original assay buffer (**Fig. 5a**). Therefore, whether these mutants exhibit greater enzyme activity than wild-type in cells may depend on the local abundance and composition of ionic species in the nucleus. Taken together, these data are consistent with the notion that DNA binding by the PWWP domain is necessary for maximal activity, and that mutations impacting this ability can modulate catalytic activity.

### DNA binding is a noncanonical PWWP domain role in cells

To evaluate whether our findings extend to DNMT3A’s native cellular context, we next considered whether PWWP domain mutations affect de novo DNA methylation in cells. We ectopically expressed mouse Dnmt3a2 variants in *Dnmt1*/*Dnmt3a*/*Dnmt3b*-triple knockout (TKO) mouse embryonic stem cells (ESCs)^47^ (**Extended Data Fig. 7a–d**) and conducted reduced representation bisulfite sequencing (RRBS)^48,49^ to measure DNA methylation. These cells are severely hypomethylated at baseline and rapidly lose methylation due to a lack of Dnmt1 maintenance, enabling us to detect where methylation is deposited with high stringency. Consistent with our in vitro results, analysis of CpG methylation genome-wide showed that Dnmt3a2 R297W (human R301W) was functionally defective compared to wild-type (**Fig. 5f**, left). These differences were even more pronounced when averaging over larger genomic bins (**Fig. 5f**, right). Nevertheless, Dnmt3a2 R297W retained partial activity in cells compared to the catalytically dead E752K mutant (human E756K). By contrast, both W326R and E338K mutants (human W330R and E342K, respectively) recovered similar amounts of methylation compared to wild-type Dnmt3a2, though we note that high replicate variability was observed for the E338K mutant. These results suggest that cellular methylation requires a minimum threshold of PWWP domain DNA affinity, although increased affinity may not necessarily confer global hyperactivity in cells.

To analyze whether these mutations affect histone targeting in cells, we performed chromatin-immunoprecipitation and sequencing (ChIP-seq) to map patterns of H3K4me3 and H3K36me2 in parent TKO ESCs. Wild-type, R297W, W326R, and E338K variants all displayed a sharp reduction in methylation at regions containing high H3K4me3 levels, as expected (**Extended Data Fig. 8a**). With regard to H3K36me2, we found that Dnmt3a2 W326R was equally able to methylate regions regardless of their H3K36me2 levels; moreover, it produced higher DNA methylation at low H3K36me2 regions than the other variants (**Fig. 5g and Extended Data Fig. 8b,c**). This is consistent with mistargeting and aberrant spreading of methylation, as well as our finding that DNMT3A2 W330R is hyperactive at baseline (**Fig. 5a**). By contrast, the wild-type, R297W, and E338K variants all displayed a positive correlation between DNA methylation and H3K36me2 levels (**Fig. 5g**), confirming that neither R297W nor E338K abrogate the PWWP domain’s canonical histone reader function. Therefore, taken together, our biochemical and genomic results show that DNA binding is distinct from the PWWP domain’s canonical H3K36me2 reader function, constituting an important noncanonical role in cellular methylation (**Fig. 6**).

**Fig. 6.**
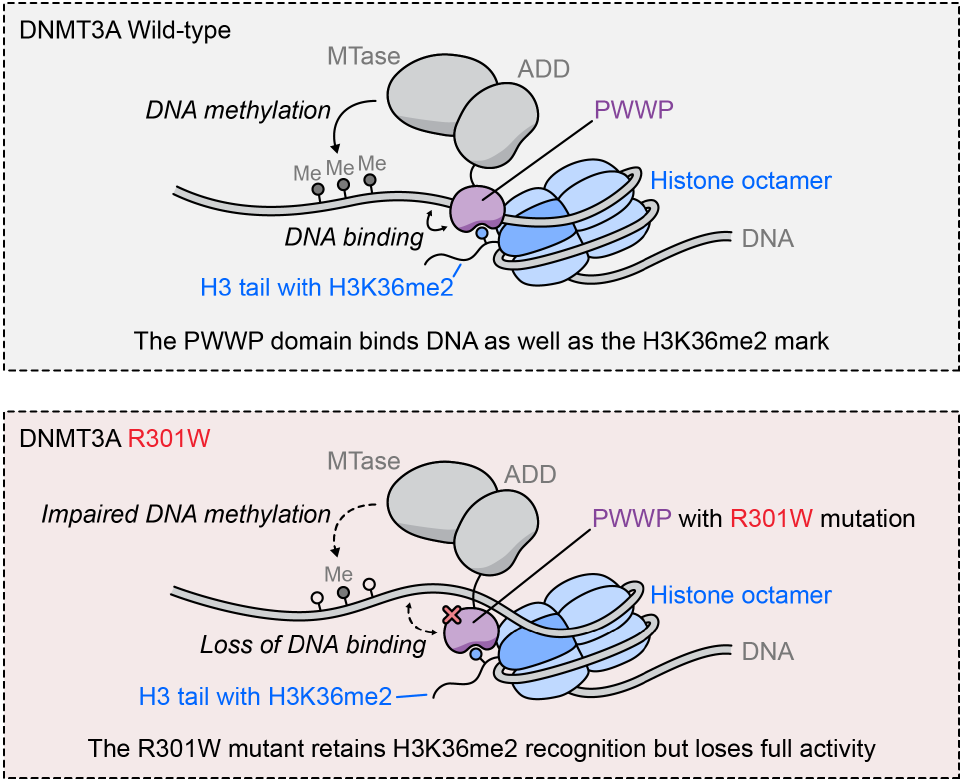
PWWP domain DNA binding is required for full activity. Model showing the role of PWWP domain DNA binding in DNMT3A methylation. Wild-type PWWP domain binds DNA in addition to its canonical role as a H3K36me2 reader (top). The R301W mutation in the PWWP domain abrogates DNA binding, leading to impaired methylation, although H3K36me2 targeting is preserved (bottom).

## Discussion

As evidenced by the growing list of developmental and hematologic disorders linked to DNMT3A, the mechanisms controlling where and when DNA methylation is installed are of paramount importance to human health. Nevertheless, fundamental questions remain unanswered regarding how DNMT3A activity is regulated, both through allosteric mechanisms intrinsic to the protein, or through interactions with factors like the nucleosome^14^. Recent studies have greatly enriched our understanding of these processes and the ways in which clinical mutations impact them^5,24,50^, but it remains challenging to probe DNMT3A function within its endogenous cellular ecosystem. Here, using a reporter for endogenous DNMT3A activity, we extend base editor screening to enable mutational scanning directly on the endogenous *DNMT3A* gene, generating key insights into the roles of the ADD and PWWP domains.

Our work identifies key features that control DNMT3A activity. We use structure-guided analysis to identify a functional hotspot at the active conformation ADD-MTase interface that is also impacted by clinical mutations^40^, providing evidence that stabilization of the active conformation, in addition to relief of autoinhibition, is necessary for full DNMT3A activity. Landmark studies have revealed that a large swath of mutations seen in patients negatively impact protein stability, in part by promoting degradation by the ubiquitin-proteasome system—particularly for mutations within the PWWP and MTase domains^5,24^. In this context, our DSF results using purified PWWP domains are intriguing as they suggest that instability can also be intrinsic to DNMT3A, for instance with the R326C clonal hematopoiesis mutant. For such mutants, cellular degradation machinery may not be necessary to mediate loss-of-function.

A central finding of our study is that DNA binding by the PWWP domain is required for optimal function. While previous studies have shown that binding of the PWWP domain to DNA contributes to substrate inhibition^45^ and can be abrogated by charge reversal mutations^44^, it is unknown whether this phenomenon contributes positively to DNMT3A function. We find that the R301W mutation impairs catalytic activity and provide evidence that loss of PWWP domain DNA binding underlies this effect (**Fig. 6**). Additionally, we observe that increased DNA binding by the E342K mutant compensates for H3K36me2 stimulation in vitro. Given the physical proximity of H3K36 to nucleosomal linker DNA, this observation could suggest that H3K36me2 stimulation normally occurs by promoting DNA binding. In this context, it is interesting to note that a recent cryo-electron microscopy study was unable to resolve the PWWP domain within full-length DNMT3A2, suggesting it is conformationally dynamic^14^. Such dynamics may underlie noncanonical pathways of allosteric feedback between the PWWP domain and other parts of DNMT3A.

Additional structural and mechanistic insight into interdomain crosstalk—particularly involving the PWWP domain—will greatly inform our understanding of DNMT3A and its role in disease. For instance, recent studies have shown that the W330R HESJAS mutation, by abrogating PWWP reader function, causes aberrant targeting to regions marked with H2AK119ub via an interaction specific to the isoform 1 N-terminus^5,50^. However, our work demonstrates that even in the context of DNMT3A2, which lacks this interaction, the W330R mutation has an effect on function. Understanding the mechanistic details of how increased PWWP domain DNA affinity promotes catalytic activity, such as for the W330R and E342K mutants, may offer insight into allosteric mechanisms controlling DNMT3A function. Such insights could open novel avenues toward therapeutically targeting DNMT3A in cancer or other diseases.

More broadly, our work offers a framework for in situ mutational scanning of other chromatin regulators. The fluorescent reporter that we adapted here has been used to characterize writer enzymes of a variety of repressive chromatin modifications^36^, and more recently has been extended to a diverse set of repressive and activating transcriptional effector domains^51^, demonstrating its versatility. By systematically probing sequence-activity relationships, our approach can yield biochemical insight even for proteins that are challenging to study in vitro. At the same time, it is not limited by target essentiality in cells or by the availability of potent drugs. We anticipate future studies employing base editor scanning will yield novel insights into chromatin regulators and the complex, intertwined mechanisms defining their activities in cells.

## Data Availability

RRBS and ChIP-seq data have been deposited to NCBI GEO. Additional data is available from the corresponding author upon request.

## Code Availability

Custom code used for analysis of base editor scanning, genomics, and other next-generation sequencing data will be made available on github prior to publication.

## Acknowledgements

We thank members of the Liau Lab for helpful discussions and comments on the manuscript, in particular A. Siegenfeld, A. Waterbury, H.S. Kwok, S. Roseman, and P. Gosavi. We thank D. Bolduc for advice regarding biochemistry experiments, Z. Niziolek and J. Nelson for assistance with FACS, A. Meissner for providing TKO ESCs, A. Mattei and E. Jung for advice regarding stem cell culture, S. Berry for computational advice, V. Baidin for advice regarding radiometric assays, and T. Haining and K. Richards-Corke for additional assistance. N.Z.L., E.M.G., and K.C.N. were supported by NSF Graduate Research Fellowships (grant no. DGE1745303). E.M.G. was also supported by the Landry Cancer Biology Fellowship. C.L. was supported by the Herchel Smith Graduate Fellowship. This research was supported by award no. 1DP2GM137494 from the National Institute of General Medical Sciences, the Damon Runyon-Rachleff Innovation Award, and startup funds from Harvard University.

## Author Contributions

N.Z.L. and B.B.L. conceived the study and wrote the manuscript with assistance from E.M.G. N.Z.L., E.M.G., and B.B.L. designed experiments and analyzed data. N.Z.L. performed base editor scanning, cell culture experiments, biochemistry experiments, coding, genomics, genomics analysis, and data visualization. E.M.G. performed conservation analysis, cell culture experiments, biochemistry experiments, coding, and genomics. K.C.N. assisted in reporter design and validation and contributed code. C.L. assisted in base editor cloning and contributed code. J.G.D. oversaw sgRNA library design and contributed the base editor vector. All authors edited and approved the manuscript. B.B.L. supervised and held overall responsibility for the study.

---

**Extended Data Fig. 1.**
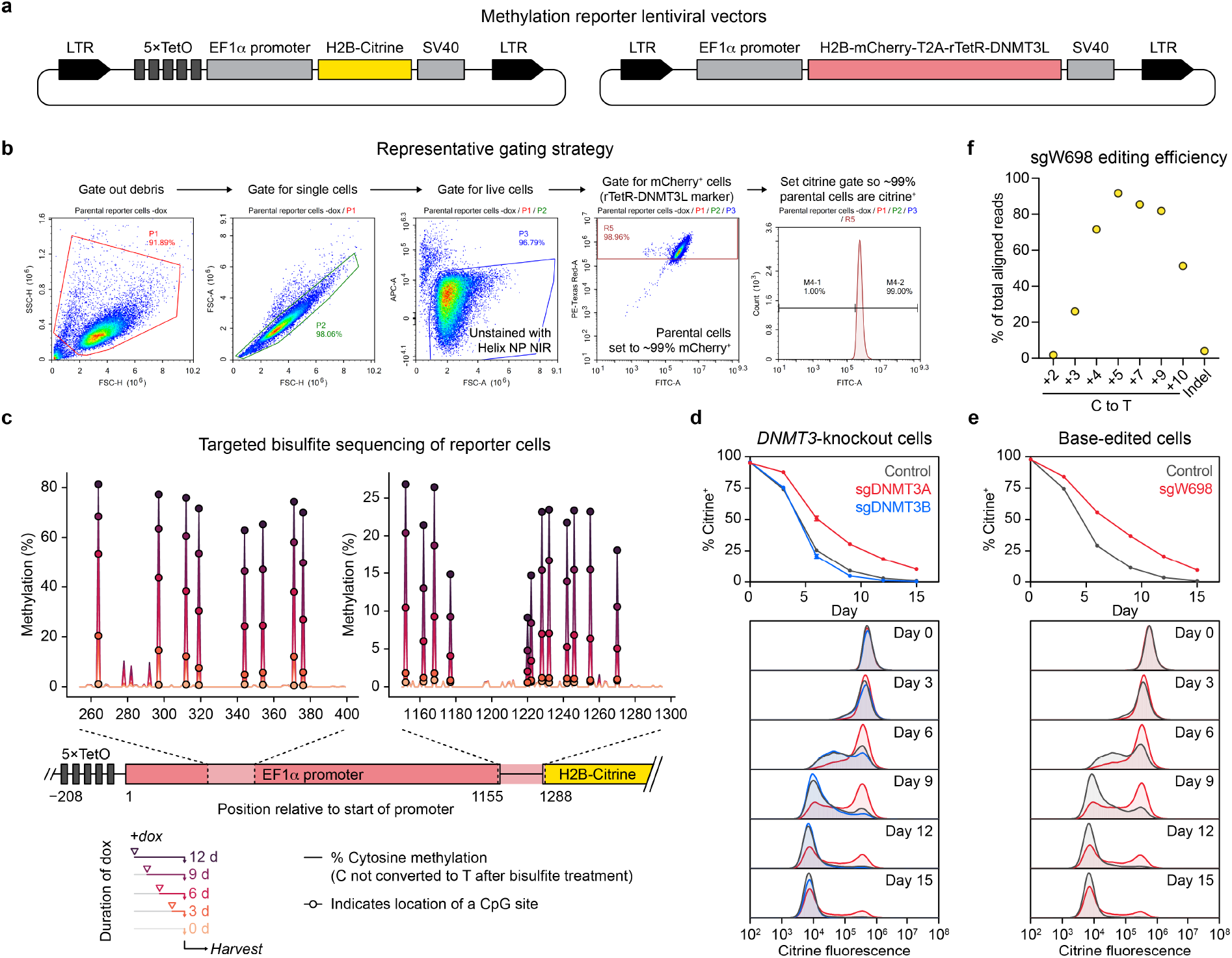
Reporter silencing depends on DNMT3A and is concomitant with DNA methylation. **a**. Schematic of lentiviral methylation reporter vectors. LTR, lentiviral long terminal repeat; TetO, tetracycline operator; SV40, simian virus 40 poly(A) sequence; rTetR, reverse Tet repressor. **b**. Representative gating scheme for flow cytometric analysis of reporter silencing assays. Helix NP NIR was used as a viability dye. Citrine fluorescence and mCherry fluorescence were monitored on the FITC and PE-Texas Red channels, respectively. **c**. Reporter methylation levels after varying duration of dox treatment measured by targeted bisulfite sequencing. In each plot, lines represent the percent cytosine methylation at each position (non-cytosine positions are set to 0). CpG sites are highlighted by dots. A schematic of the reporter is shown below, indicating the location of each sequenced amplicon. This experiment was performed once, with n = 1. **d**. Full timecourse for *DNMT3*-knockout silencing experiment shown in Fig. 1f (top), with representative histograms of citrine fluorescence from each day (bottom). **e**. Full timecourse for sgW698 silencing experiment shown in Fig. 1h (top), with representative histograms of citrine fluorescence from each day (bottom). **f**. Deep sequencing of cells edited with sgW698 after 9 d of dox treatment, and sorted for citrine^+^ cells. Plot shows the percentage of aligned reads with C to T base edits at each indicated protospacer position, or the percentage of aligned reads with indels. Sequencing was performed once. For **d** and **e**, data and error bars (where larger than data point) are mean ± SD of n = 3 replicates, and one of two independent experiments is shown.

**Extended Data Fig. 2.**
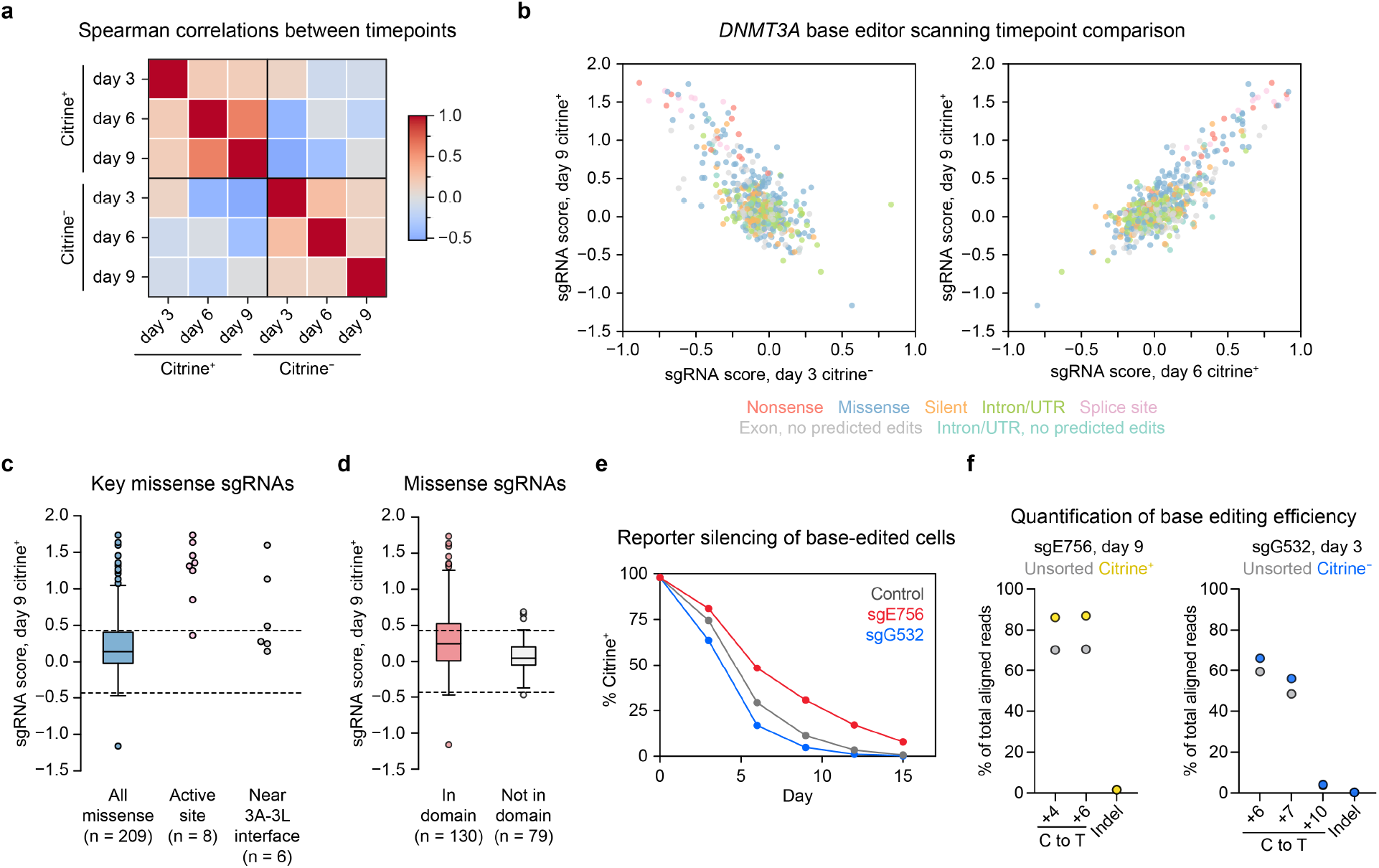
Analysis and validation of DNMT3A base editor scanning results. (**a–d**) *DNMT3A* base editor scanning results. Data are the average of n = 3 replicates. For **c** and **d**, dotted lines indicate ±2 SD of intergenic control sgRNAs, and boxplot components are as follows: center line, median; box, interquartile range; whiskers, up to 1.5 × interquartile range per the Tukey method. **a**. Heatmap depicting Spearman correlations between sgRNA scores obtained at different timepoints, and for citrine^+^ or citrine^−^ cells, in the base editor scanning experiment. **b**. Scatterplots showing the negative correlation between sgRNA scores for the day 9 citrine^+^ and day 3 citrine^−^ samples (left), and the positive correlation between sgRNA scores for the day 9 citrine^+^ and day 6 citrine^+^ samples (right). **c**. sgRNA scores at day 9 in citrine^+^ cells for missense sgRNAs targeting the active site or residues near the DNMT3A-DNMT3L interface, compared to all missense sgRNAs. **d**. sgRNA scores at day 9 in citrine^+^ cells comparing missense sgRNAs targeting the PWWP, ADD, or MTase domains (red) or targeting a non-domain region of DNMT3A (light gray). **e**. Full timecourse for the sgE756 and sgG532 silencing experiment shown in Fig. 2e. Data are mean ± SD of n = 3 replicates, and one of two independent experiments is shown. **f**. Deep sequencing of cells edited with sgE756 after 9 d of dox treatment (left) or with sgG532 after 3 d of dox treatment (right), either unsorted or sorted as indicated. Plots show the percentage of aligned reads with C to T base edits at each indicated protospacer position, or the percentage of aligned reads with indels. Sequencing was performed once.

**Extended Data Fig. 3.**
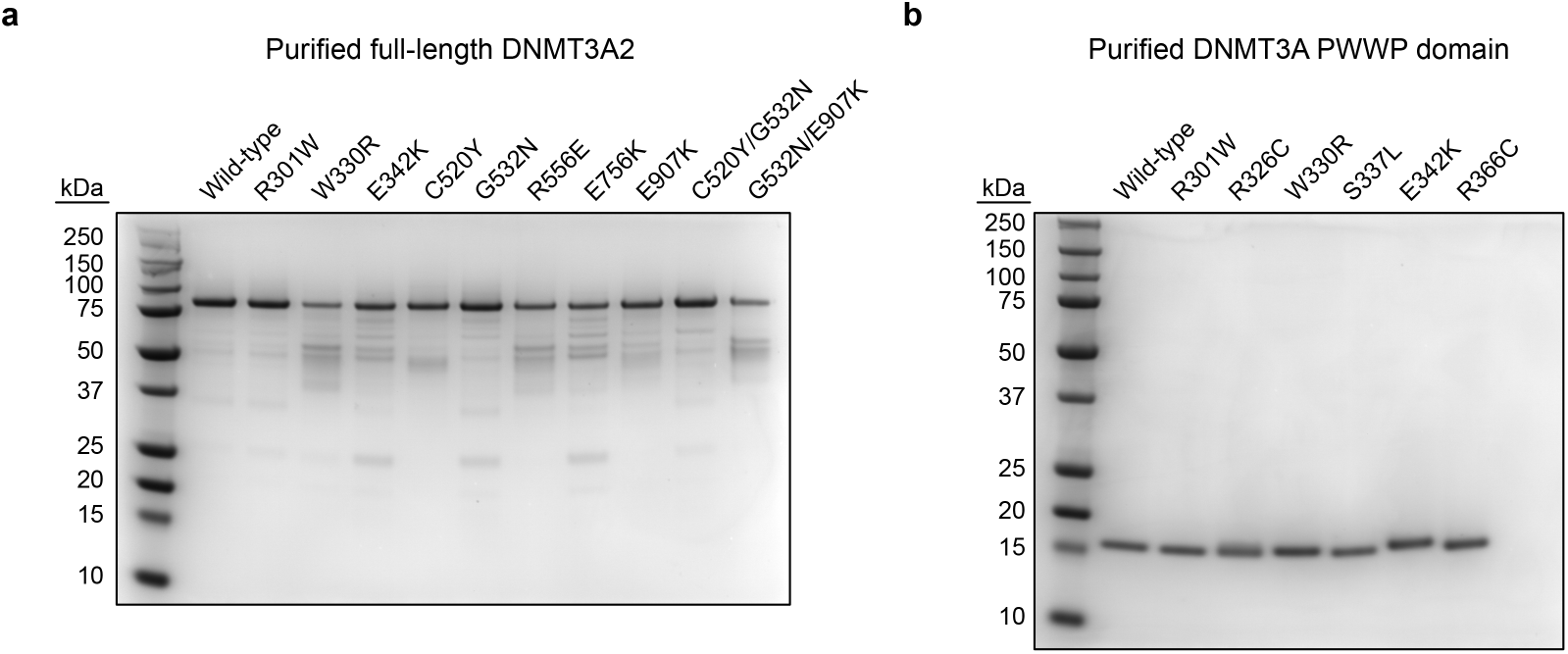
SDS-PAGE of purified proteins. **a–b**. Purified (**a**) full-length DNMT3A2 (80 kDa, residues 224–912 with N-terminal 6×His tag) and (**b**) PWWP domains (17 kDa, residues 89–238, untagged), electrophoresed on Novex 10% acrylamide tricine gels (Thermo Fisher Scientific) and visualized by Coomassie staining. 1 μg protein was loaded in each lane. Protein purifications were generally performed once, although wild-type DNMT3A2 was purified more than once and verified to have comparable activity across purifications.

**Extended Data Fig. 4.**
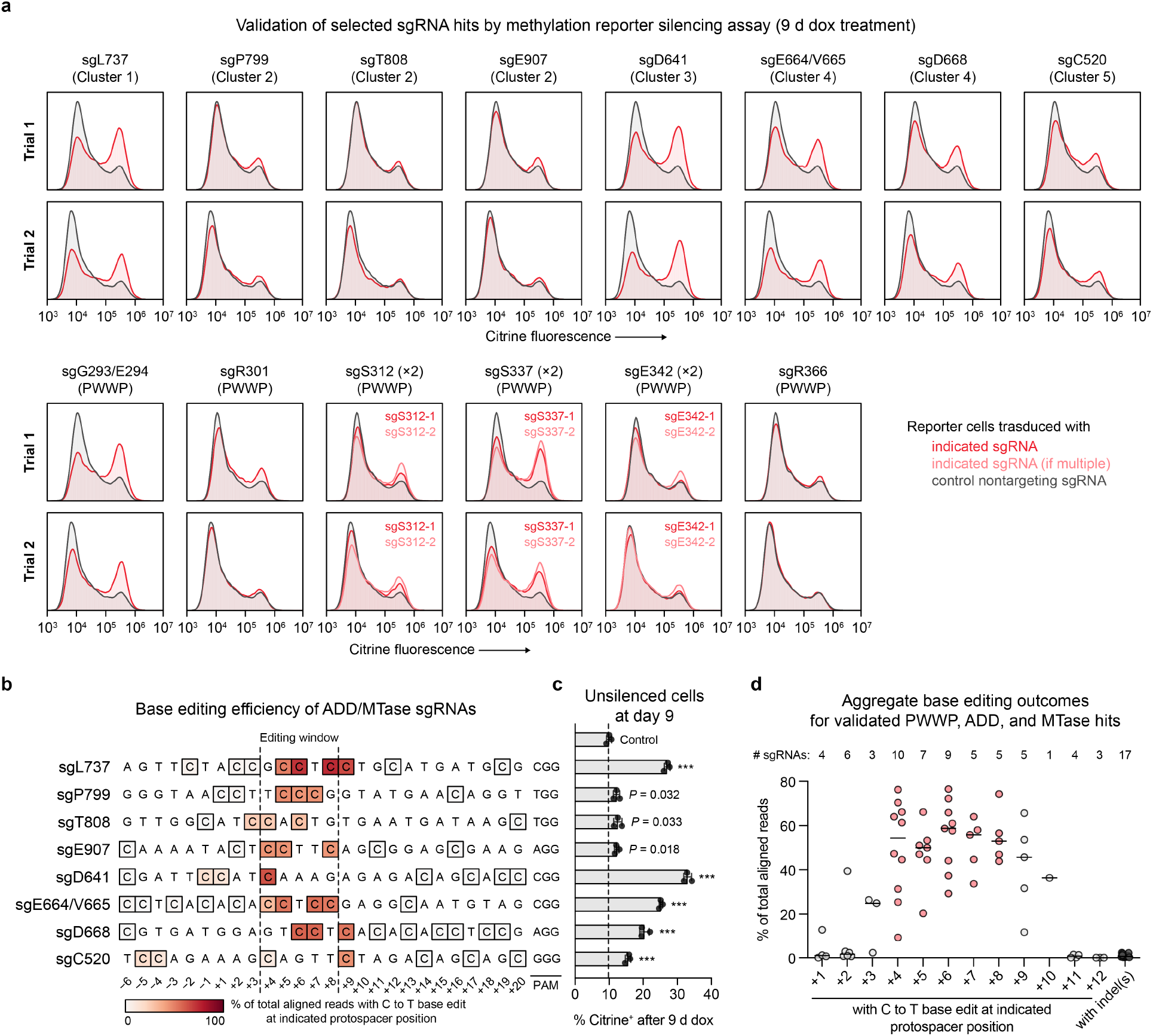
Validation of individual sgRNA hits. **a**. Citrine fluorescence of base-edited cells after 9 d of dox treatment for 17 sgRNAs targeting *DNMT3A* (red or light red). Histograms are representative of n = 3 replicates. Top and bottom plots show data from the two experimental trials (independent transductions). The citrine fluorescence histogram of control cells treated with dox in parallel (gray) is overlaid in each plot. Control data shown are identical for samples analyzed in the same experiment. Data shown in Fig. 4d corresponds to trial 2 shown here. **b**. Next-generation sequencing analysis of base editing efficiency at each editable C within the protospacer of the indicated sgRNAs. **c**. Flow cytometric quantification of cells remaining citrine^+^ after 9 d of dox treatment. Data correspond to those shown in **a**, trial 2, and are mean ± SD of n = 3 replicates. *P* values (***, *P* ≤ 0.001; ns, not significant) were calculated through two-tailed unpaired *t* tests comparing each sgRNA to control. **d**. Aggregate base editing outcomes for the 17 sgRNAs presented in **a** (includes PWWP, ADD, and MTase hit sgRNAs). The efficiency of base editing is plotted at each protospacer position for all sgRNAs containing a C at that position. Horizontal lines indicate the median at each position. The number of sgRNAs with a C at each position is printed above. The indel frequencies for all sgRNAs is shown to the right in dark gray. Protospacer positions within the editing window (+4 to +8) are highlighted in red. Genotyping was performed once for each sgRNA.

**Extended Data Fig. 5.**
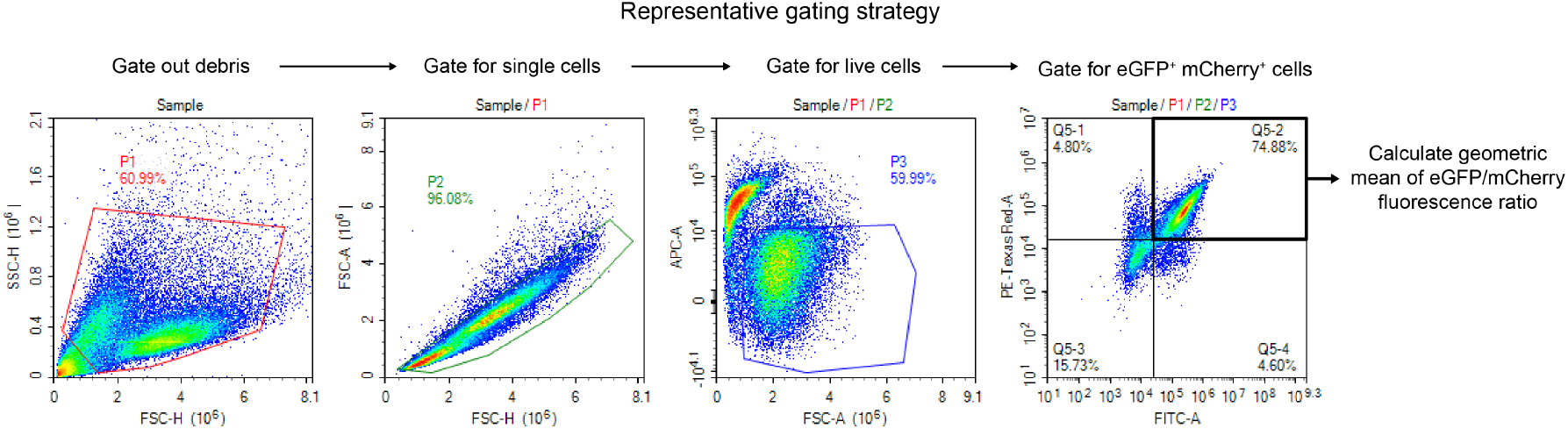
Stability reporter gating strategy. Representative gating strategy used for stability reporter assay measurements. Quadrant gate in mCherry vs. eGFP plot was set so that wild-type parent K562 cells were mCherry/eGFP double-negative.

**Extended Data Fig. 6.**
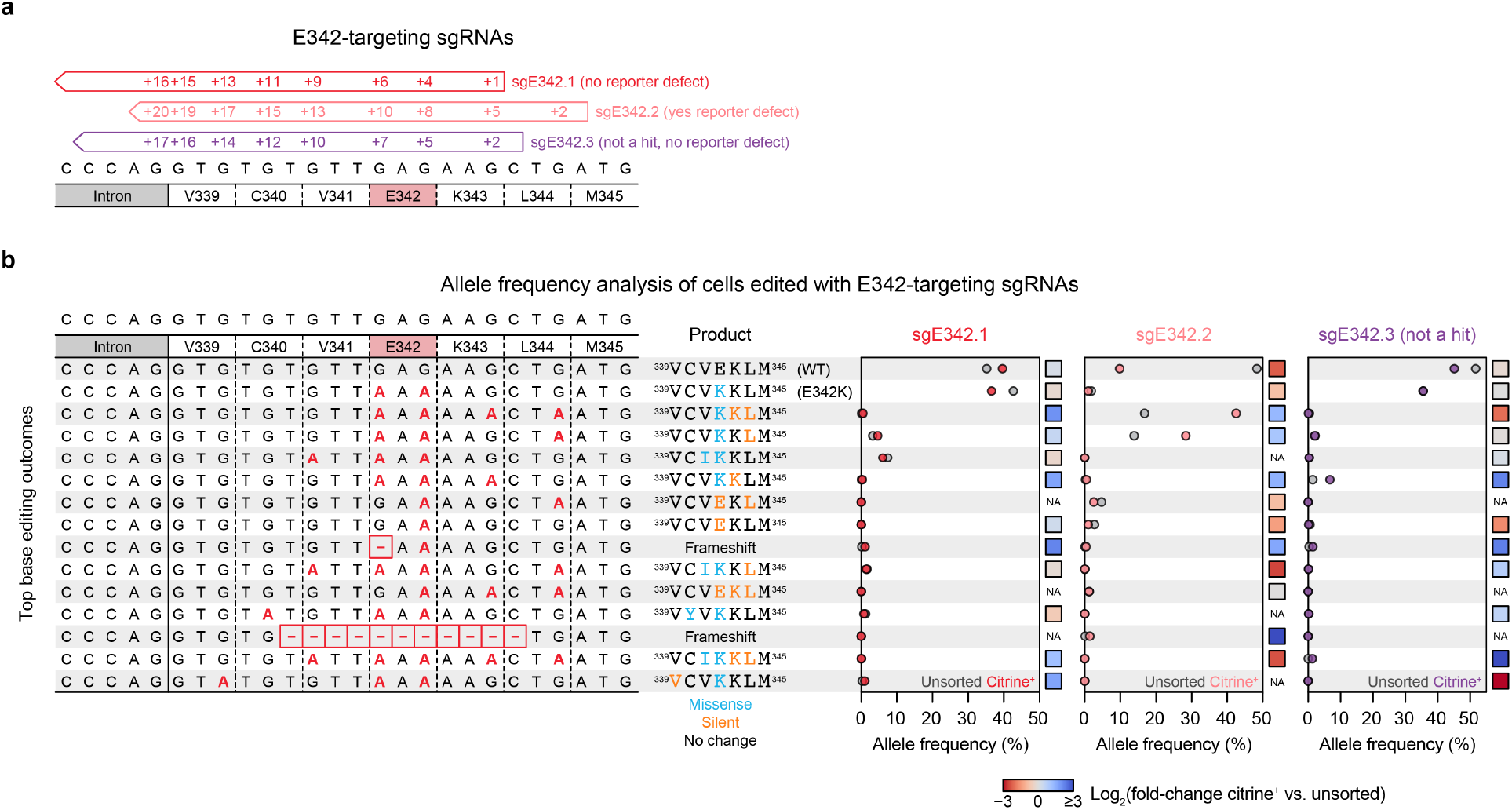
Analysis of base editing outcomes for three sgRNAs targeting the E342 codon of DNMT3A. **a**. Schematic of sgRNAs targeting the E342 codon. sgE342-1 (red) and sgE342-2 (light red) correspond to the screening hits presented in Figures 4 and S3. sgE342-3 (purple) is an additional library sgRNA that did not score as a hit in the screen but also targets the E342 codon. **b**. Allele frequencies in base editor-treated reporter cells after 15 days of dox treatment, comparing citrine^+^ cells to unsorted cells. Genomic DNA was harvested using QuickExtract DNA extraction solution (Lucigen) and libraries were prepared and deep sequenced as for other genotyping experiments. Each row represents an allele (for the purposes of this analysis, alleles were merged if they were identical within the region depicted here). All alleles having at least 1% allele frequency in at least one sample are depicted. Left, nucleotide sequence of each allele, with C to T base edits shown in red (these appear as G to A because the protospacers are along the opposite strand) and dashes representing deletions. Middle, amino acid sequence corresponding to the translation of the region shown, with missense and silent mutations colored blue and orange, respectively. Right, allele frequency in unsorted or citrine^+^ cells for each sgRNA. Colored dots, citrine^+^ cells; gray dots, unsorted cells. Colored squares to the right of each plot indicate the log_2_(fold-change in allele frequency in citrine^+^ vs. unsorted cells). NA indicates undefined log_2_(fold-changes) where one or both of the allele frequencies is zero. This experiment was conducted once (n = 1).

**Extended Data Fig. 7.**
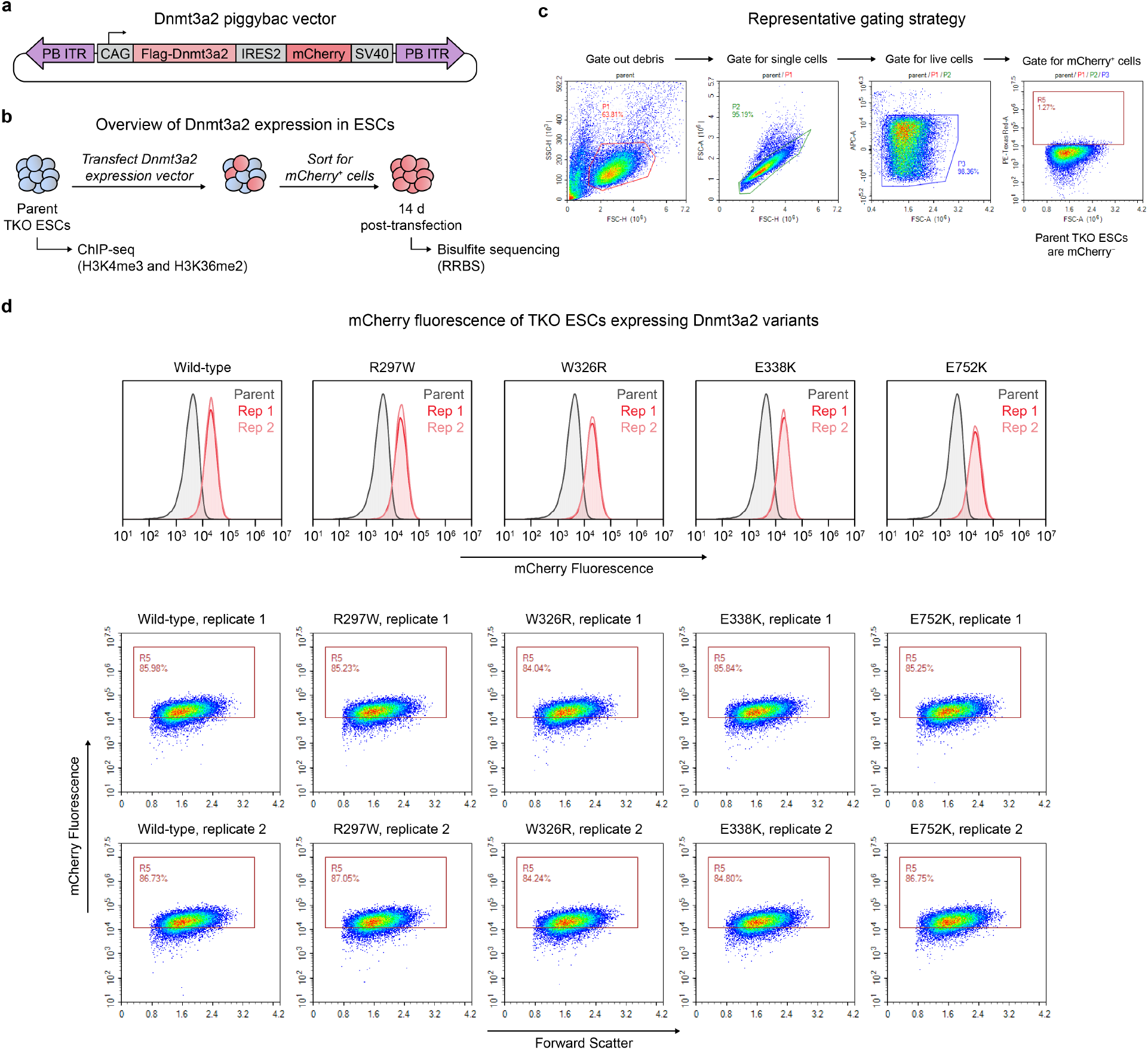
Generation of Dnmt3a2-complemented TKO ESCs. **a**. Schematic of piggybac (PB) vector used for ectopic Dnmt3a2 expression in TKO ESCs. ITR, inverted terminal repeat. **b**. Overview of Dnmt3a2 complementation experiment. **c**. Representative gating strategy for flow cytometric analysis and FACS of TKO ESCs. **d**. Flow cytometric analysis of mCherry fluorescence in ESCs transposed with the Dnmt3a2 expression vector. Top, histograms of mCherry fluorescence. Parent TKO ESCs are shown in gray (same data for all plots). Bottom, mCherry fluorescence versus forward scatter showing the percent of gated mCherry^+^ cells. Replicates 1 and 2 correspond to separately transfected cells.

**Extended Data Fig. 8.**
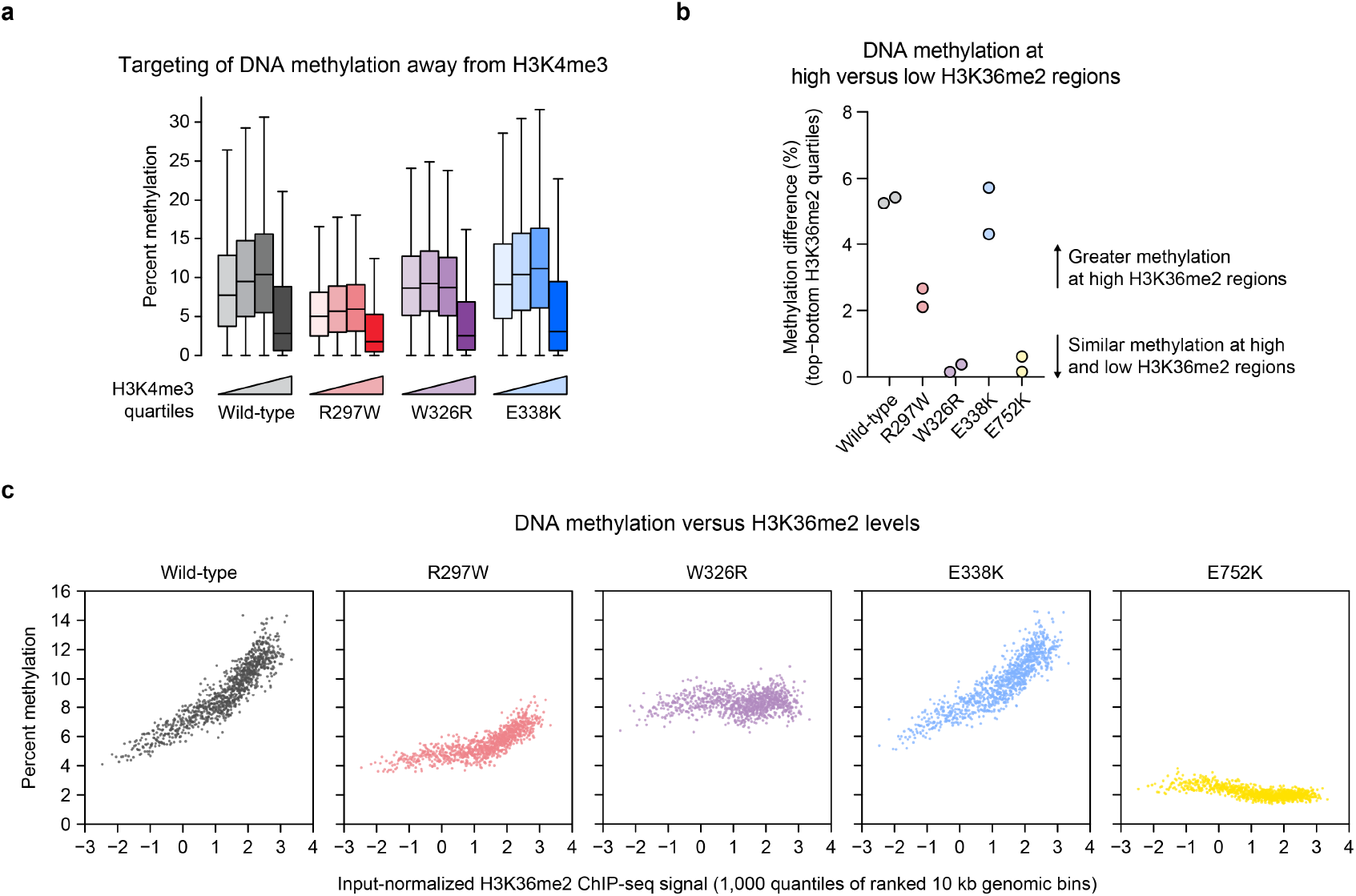
Analysis of de novo DNA methylation at H3K4me3- and H3K36me2-marked regions. **a**. CpG methylation within 10 kb genomic bins ranked into quartiles based on normalized H3K4me3 ChIP-seq signal (n = 93,516 bins total, 23,379 bins per quartile). **b**. Difference in methylation between top and bottom H3K36me2 quartiles for each sample. 10 kb genomic bins (n = 93,516 total) were grouped into quartiles, and the average methylation in the median bins from the top and bottom quartiles were compared. Biological replicates are shown separately. **c**. CpG methylation within 10 kb genomic bins ranked into quantiles (n = 1,000 quantiles) based on normalized H3K36me2 ChIP-seq signal. The average bin methylation for each quantile is plotted against the H3K36me2 signal. For **a–c**, only CpGs with 5× coverage across all samples were considered. Methylation values in **a** and **c** represent the average of two biological replicates.

## Methods

### Cell culture

K562 was obtained from ATCC. 293FT was obtained from Thermo Fisher Scientific. HEK293T was a gift from B.E. Bernstein (Massachusetts General Hospital). *Dnmt1*/*Dnmt3a*/*Dnmt3b*-triple knockout (TKO) mouse embryonic stem cells (ESCs) were a gift from A. Meissner (Max Planck Institute). CD-1 mouse embryonic fibroblasts (MEFs) were obtained from Lonza (Cat#M-FB-481) and inactivated with 10 μg ml^−1^ mitomycin C (Sigma-Aldrich) for use as feeder cells. All cell lines were cultured in a humidified 5% CO_2_ incubator at 37 °C and were tested for mycoplasma. All media were supplemented with 100 U ml^−1^ penicillin and 100 μg ml^−1^ streptomycin (Gibco). Fetal bovine serum (FBS) was obtained from Peak Serum except for ESC medium. K562 cells were cultured in RPMI-1640 (Gibco) with 10% FBS. HEK293T cells were cultured in DMEM (Gibco) with 10% FBS. 293FT cells were cultured in DMEM with 10% FBS, 2 mM GlutaMAX (Gibco), and 1× MEM non-essential amino acids (Gibco). ESCs were cultured in KnockOut DMEM (Gibco) with 15% FBS (Gibco, heat-inactivated), 2 mM GlutaMAX, 1× MEM non-essential amino acids, 10^3^ U ml^−1^ ESGRO leukemia inhibitory factor (Millipore), and 55 μM 2-mercaptoethanol (Gibco). ESCs were cultured on a layer of MEF feeders plated on 0.2% gelatin-coated vessels. ESC medium was changed daily, and feeders were depleted prior to sample harvest.

### Lentiviral cell line generation

Lentiviral transduction was used to establish methylation reporter and stability reporter cell lines and introduce sgRNAs for base editing and CRISPR knockout. To produce lentivirus, transfer plasmid was co-transfected with *GAG/POL* and *VSVG* plasmids into HEK293T or 293FT cells using FuGENE HD (Promega, 3.33:1 FuGENE:DNA) and the medium was replaced 6–8 h after transfection. 48–60 h later, the medium was collected, passed through a 0.45 μm filter, snap-frozen, and stored at –80 °C until use. K562 cells were transduced by spinfection at 1,800*g* for 90 min at 37 °C with 12 μg ml^−1^ polybrene (Santa Cruz Biotechnology) and the appropriate lentivirus. Successfully transduced cells were selected using 2 μg ml^−1^ puromycin (Gibco) or through fluorescence-activated cell sorting (FACS). A clonal K562 methylation reporter cell line was first derived and used in all subsequent base editor scanning and sgRNA validation experiments. Bulk selected cells were used for additional experiments.

### Plasmid construction

Plasmids for sgRNA expression were cloned by annealing synthetic oligonucleotides (Sigma-Aldrich) and ligating them into the appropriate vector, either lentiCRISPR v2, a gift from F. Zhang (Addgene #52961, for Cas9 knockout), or pRDA_254, a modified version of pRDA_256 (ref. ^32^) expressing BE3.9max-P2A-PuroR but lacking the 10× guide capture sequence (for base editing). All other plasmids were cloned by Gibson Assembly using NEBuilder HiFi (New England Biolabs). Mutations were introduced using primers containing the desired mutations. The following cloning strains were used: NEB Stable (lentiviral and piggybac vectors) and NEB 5-alpha (all other plasmids) (New England Biolabs). For cloning of base editor constructs, bacterial cultures were grown at 30 °C to avoid plasmid recombination. Final constructs were validated by Sanger sequencing (Quintara Biosciences).

In general, all plasmids used in this study for expression of DNMT3A encode isoform 2 (human DNMT3A2: residues 224–912; mouse Dnmt3a2: residues 220–908), unless otherwise indicated. The coding sequences of human DNMT3A and DNMT3L for mammalian expression were amplified from pcDNA3/Myc-DNMT3A and pcDNA3/Myc-DNMT3L, respectively, gifts from A. Riggs (Addgene #35521, 35523). The methylation reporter was constructed using sequences derived from PhiC31-Neo-ins-5xTetO-pEF-H2B-Citrine-ins and pEX1-pEF-H2B-mCherry-T2A-rTetR-KRAB, gifts from M. Elowitz (Addgene #78099, 78348). Specifically, two cassettes, a reporter (5xTetO-pEF-H2B-Citrine-SV40) and silencer (pEF-H2B-mCherry-T2A-rTetR-DNMT3L-SV40), were each cloned into LT3REVIR, a gift from J. Zuber (Addgene #111176). To clone stability reporter constructs, DNMT3A2 variants were inserted into a modified version of Cilantro2, a gift from B. Ebert (Addgene #74450). The pET28b-His_6_-DNMT3A2 bacterial expression plasmid was a gift from D.M. Bolduc (H3 Biomedicine). For bacterial expression of isolated PWWP domains, residues 278–427 of DNMT3A were cloned into pET28b with an N-terminal His_6_-MBP tag. This was designed so that TEV protease cleavage recovers the PWWP domain with no appended residues. The coding sequence of mouse Dnmt3a2 (NM_007872.4) was synthesized in two pieces by Twist Bioscience and cloned as a CAG-Flag-Dnmt3a2-Ires2-mCherry-SV40 cassette into the Piggybac vector pSLQ2817, a gift from S. Qi (Addgene #84239).

### Reporter silencing assays

Reporter cells were plated in triplicate at 1 × 10^5^ cells ml^−1^ in media with or without 100 ng ml^−1^ doxycycline (dox) (Sigma-Aldrich). Every 3 d, samples were removed for analysis and cells were passaged by dilution into fresh media with or without dox. For washout, cells were washed once with phosphate-buffered saline (PBS) and resuspended in fresh media without dox. For flow cytometry, Helix NP NIR viability dye (BioLegend) was added and data were collected on a NovoCyte 3000RYB and analyzed using NovoExpress (ACEA). Gates were set so that ∼99% of mCherry^+^ reporter cells, cultured in parallel in the absence of dox, were citrine^+^ (see **Extended Data Fig. 1b** for a representative gating scheme). Reporter silencing assays were each performed twice using independently transduced cells, with similar results.

### Genotyping of base-edited cells

Genomic DNA was purified using the QIAamp DNA Blood Mini or UCP DNA Micro kits (Qiagen), unless otherwise specified. 100 ng DNA was used as input in a first round of PCR (25–27 cycles, Q5 hot start high-fidelity DNA polymerase, New England Biolabs) to amplify the genomic loci of interest and attach common overhangs. PCR products were purified by a 1.5× SPRI clean-up (Mag-Bind Total Pure NGS beads, Omega Bio-Tk), and 5 ng of each was amplified in a second round of PCR (8 cycles) to attach barcoded adapters. The final amplicons were pooled, purified by gel extraction (Zymo), and sequenced on an Illumina MiSeq. Data were processed using CRISPResso2 (ref. ^52^) in batch mode using the following parameters: --quantification_window_size 20 --quantification_window_center -10 -- plot_window_size 20 --exclude_bp_from_left 0 --exclude_bp_from_right 0 -- min_average_read_quality 30 --n_processes 12 --base_editor_output. Custom python scripts were used to extract data from the output and construct allele tables.

### Reporter bisulfite sequencing

Reporter cells were treated with 100 ng ml^−1^ dox starting at staggered timepoints, so that all samples were harvested simultaneously. Genomic DNA was extracted and subjected to bisulfite conversion using the EZ DNA Methylation kit (Zymo). Sequencing libraries were prepared following the same protocol used for genotyping, except EpiMark hot start *Taq* DNA polymerase (New England Biolabs) was used. Bisulfite-compatible primers used to amplify regions of the methylation reporter were a gift from D.M. Bolduc (H3 Biomedicine) or were designed using MethPrimer^53^. Data were processed using CRISPResso2 (ref. ^52^) with default settings. A custom python script was used to compute the percent methylation at each amplicon position containing a bisulfite-convertible base (C or G, depending on the direction of sequencing) in the reference sequence. Percent methylation was calculated as (reads with bisulfite-unconverted base, C or G) / (reads with bisulfite-converted or -unconverted base, either C/T or G/A) × 100. Elsewhere, this value was set to zero.

### Immunoblotting

Cells were washed three times with PBS and lysed by incubation for 30 min on ice in RIPA lysis buffer (Boston BioProducts) supplemented with 1× Halt Protease Inhibitor Cocktail and 5 mM EDTA (Thermo Fisher Scientific). Lysate was clarified by centrifugation and the total protein concentration was measured with the BCA Protein Assay (Thermo Fisher Scientific) to enable input standardization. The insoluble lysate fraction was added back to the clarified lysate prior to preparing samples for electrophoresis. Immunoblotting was performed according to standard procedures using the following primary antibodies: DNMT3A (#D2H4B, Cell Signaling Technology, 1:5,000), DNMT3B (#D7070, Cell Signaling Technology, 1:1,000), GAPDH (#sc-47724, Santa Cruz Biotechnology, 1:2,000). Immunoblots were visualized by film exposure using horseradish peroxidase-conjugated secondary antibodies and SuperSignal West Femto (DNMT3A) or SuperSignal West Pico PLUS (other proteins) chemiluminescent substrates (Thermo Fisher Scientific).

### Library cloning and base editor scanning

The sgRNA library was designed as described previously^32^ to include all sgRNAs with an NGG protospacer-adjacent motif targeting exonic and flanking intronic regions of *DNMT3A* isoform 2 (NM_153759.3, ENST00000380746), excluding promiscuous sgRNAs and those with TTTT sequences. Negative (nontargeting, intergenic) and positive (essential splice site) controls were included. The library was ordered as an oligonucleotide pool from Twist Biosciences and cloned into pRDA_254 according to previously described pooled cloning workflows^54,55^. Lentivirus was produced from the plasmid library and titered by measuring cell counts after transduction and puromycin selection. 21 × 10^6^ reporter cells were transduced with library virus at a multiplicity of infection less than 0.3 and selected with puromycin for 7 d. Cells were then expanded and split into three replicate subcultures and treated with 100 ng ml^−1^ dox. Cultures were passaged as necessary. 3 d, 6 d, and 9 d after starting dox treatment, cells were sorted on a FACSAria II (BD), collecting citrine^+^, citrine^−^, and unsorted (all mCherry^+^) cells. Genomic DNA was isolated using the QIAamp DNA Blood Mini kit and sgRNA sequences were amplified by PCR using barcoded primers, purified by gel extraction, and sequenced on an Illumina MiSeq as previously described^25,26^. At all steps, sufficient coverage of the sgRNA library was maintained in accordance with published recommendations^55^.

### Base editor scanning data analysis

All data processing and analysis were performed using Python version 3.7.3 (www.python.org). sgRNA scores were calculated as previously described^25,26^. Briefly, sequencing reads that perfectly match each library sgRNA were counted and quantified as reads per million total reads, increased by a pseudocount of 1, log_2_-transformed, and normalized by subtracting the corresponding value in the plasmid library. One sgRNA was excluded from this and all further analysis because zero reads were detected at the day 0 timepoint. Replicates were averaged and the resulting sgRNA abundances in sorted cells were normalized by subtracting the corresponding values for matched timepoint unsorted cells. The mean value for intergenic negative controls was subtracted from each library sgRNA to calculate the final sgRNA score. sgRNAs with scores over 2 standard deviations above or below the mean of intergenic negative controls were considered “enriched” or “depleted” hits, respectively.

Each sgRNA targeting *DNMT3A* was classified based on its expected editing outcome using a custom python script. Any C within the editing window (defined as protospacer +4 to +8 positions, where +1 is PAM-distal) was assumed to be converted to T, regardless of sequence context. Based on this, sgRNAs were placed in one of 7 mutually exclusive classes, listed in order of assignment priority: (1) nonsense; (2) splice site; (3) missense; (4) silent; (5) exon, no predicted edits (no C within window); (6) intron/UTR; (7) intron/UTR, no predicted edits.

### Proximity-weighted enrichment score (PWES) analysis

The PWES for a given pair of sgRNAs *i* and *j* was calculated as previously described^25^ using the following formula based on the CLUMPS algorithm^56^:

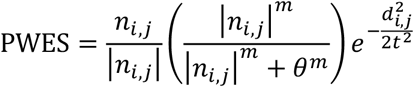

where *n*_*i,j*_ is the sum of the day 9 citrine^+^ sgRNA scores for *i* and *j, d*_*i,j*_ is the Euclidean distance between their targeted residues, and *m* = 2, *θ* = 0.8, *t* = 16. For this analysis, only missense sgRNAs editing residues resolved in the structures of active (PDB: 4U7T) and autoinhibited (PDB: 4U7P) DNMT3A were considered. sgRNAs predicted to mutate two residues were assigned to the even-numbered residue. Hierarchical clustering was performed based on the active conformation scores (PWES_4U7T_), as described previously^25^. ΔPWES was defined as ΔPWES = |PWES_4U7T_| – |PWES_4U7P_|. For each sgRNA, this metric was summed over all other sgRNAs to construct the summed ΔPWES (equivalent to a column sum of the ΔPWES matrix, excluding the diagonal).

### Conservation analysis

The protein sequence of full-length human DNMT3A (uniprot Q9Y6K1) was used as a query for five iterations of jackhmmer^57^ to search in the uniref database^58^ for sequences with homology to DNMT3A. hmm searches were performed within these sequences using the hmm profiles from pfam^59^ for the PWWP domain (version 17), ADD domain (version 1) and MTase domain (version 17), identifying subsets of these sequences containing each domain. These were intersected to generate a final list of sequences containing homology to all three DNMT3A domains. Then, the same PWWP hmm profile was used to search in uniref to gather sequences from a wide diversity of proteins that contain PWWP domains. Clustal omega^60^ was used to align both sets of sequences, and the relative level of conservation at each position in the DNMT3A PWWP domain was assessed from each alignment and compared.

### Stability reporter assay

Wild-type K562 cells were transduced with the appropriate lentivirus, selected with puromycin, and analyzed by flow cytometry. After gating for eGFP^+^ mCherry^+^ cells (see **Extended Data Fig. 5** for gating scheme), the ratio of the eGFP geometric mean fluorescence divided by the mCherry geometric mean fluorescence was calculated. All measurements were normalized to that of wild-type DNMT3A2 measured in parallel. Each mutant was assessed by two measurements of each of two independently transduced cell lines.

### Protein expression and purification

Full-length DNMT3A2 was expressed recombinantly in Rosetta2(DE3)pLysS cells (Novagen). Freshly transformed cells were grown in LB supplemented with 50 μg ml^−1^ kanamycin and 50 μg ml^−1^ chloramphenicol at 37 °C to an optical density at 600 nm of 0.6–0.8. Cells were then cooled on ice and induced with 1 mM isopropyl-Δ-D-thiogalactoside (IPTG) (Research Products International) at 16 °C overnight. Harvested cells were stored at –80 °C until use. Purification was performed according to published protocol^16^ with some modifications. Cells were resuspended in lysis buffer (base buffer (50 mM Tris-HCl, pH 8.0 cold, 300 mM NaCl, 1 mM TCEP) with 10 mM imidazole and 0.1% Triton X-100) and sonicated using a Branson sonifier. The lysate was clarified by centrifugation and further diluted with lysis buffer to a final volume of 175 ml per liter of expression culture. Diluted lysate was incubated with 1.25 ml His60 Ni Superflow affinity resin (Takara) per liter of expression culture. The resin was washed with base buffer containing a stepwise gradient of 20 mM, 50 mM, and 100 mM imidazole, followed by elution using base buffer with 200 mM imidazole. The eluate was exchanged into storage buffer (base buffer with no imidazole) using an Econo-Pac 10DG desalting column (Bio-Rad). Concentration was performed using Amicon Ultra 30 kDa centrifugal filters (Millipore) with resuspension of the sample in between 2 min spins at 2,500*g*. Frequent resuspension was necessary to prevent over-concentration, which resulted in reduced activity and/or precipitation of the protein.

DNMT3A PWWP domain was expressed in BL21(DE3) cells (New England Biolabs) using the same protocol as above, except cells were grown in TB supplemented with 50 μg ml^−1^ kanamycin and induced with 0.2 mM IPTG. Cells were resuspended and sonicated in lysis buffer (PWWP base buffer (20 mM Tris-HCl, pH 7.5 cold, 300 mM NaCl, 5% glycerol) with 10 mM imidazole and cOmplete, EDTA-free protease inhibitor (Roche)). Clarified lysate was subjected to Ni affinity purification, eluting with PWWP base buffer with 250 mM imidazole. The eluate was exchanged into cleavage buffer (50 mM Tris-HCl, pH 7.5 cold, 75 mM NaCl, 5% glycerol, 1 mM dithiothreitol (DTT), 0.5 mM EDTA) and incubated with TEV protease overnight at 4 °C. The cleaved His_6_-MBP tag was removed by incubation with Ni resin. The cleaved PWWP domain was subsequently purified using a HiTrap Heparin HP (Cytiva) to remove nucleic acid contamination (20 mM sodium phosphate, pH 7.5, 5% glycerol, 0–1.5 M NaCl gradient). Finally, protein was polished on a Superdex 200 Increase 10/300 GL column (Cytiva), eluting in storage buffer (20 mM Tris-HCl, pH 7.5 cold, 150 mM NaCl, 5% glycerol). We note that the S337L mutant was not purified by Heparin column.

Purified proteins were quantified through their absorbances at 280 nm using the calculated extinction coefficients from Expasy ProtParam. Proteins were flash frozen in liquid N_2_ and stored at –80 °C until use.

### DNMT3A activity assays

DNMT3A activity was measured using a previously described protocol^16^ with some modifications. Reactions were conducted in 50 μl volume using 10 μM base pairs poly(dI-dC) (Sigma-Aldrich); 0.5 μM adenosyl-L-methionine, *S*-[methyl-^3^H] (^3^H-SAM) (PerkinElmer, 18.0 Ci/mmol); and 0.1 μM purified DNMT3A2 in assay buffer (50 mM sodium phosphate, pH 7.5, 20 mM NaCl, 1 mM TCEP, 0.1 mg ml^−1^ bovine serum albumin (BSA), 0.01% Triton X-100). Reactions were initiated by addition of ^3^H-SAM, incubated at room temperature for 2 h, and then terminated by addition of ∼2,000-fold molar excess of nonradioactive SAM (New England Biolabs) in 300 μl assay buffer. 40 μl DEAE sepharose fast flow resin (Cytiva) was added, and samples were rotated at room temperature to allow radioactively labeled DNA to bind the resin. Resin was recovered using a Pierce spin column (Thermo Fisher Scientific), washed with 2 × 250 μl wash buffer (50 mM sodium phosphate, pH 7.5, 20 mM NaCl, 1 mM TCEP), resuspended in 200 μl H_2_O, and mixed with 4 ml Ultima Gold scintillation cocktail (PerkinElmer). Each sample was counted for 5 min on a Beckman LS 6000SC liquid scintillation counter.

For histone stimulation experiments, 1 μM peptide was added prior to enzyme. The following peptides were used: H3K4me0 (ARTKQTARKSTG-NH_2_, Biomatik), H3K36me0 and H3K36me2 (histone H3 (21–44)-GK(biotin), Anaspec, Cat#AS-64440 and AS-64442).

For the NaCl titration experiment, a modified buffer was used during the reaction incubation (20 mM HEPES, pH 7.5, 1 mM TCEP-HCl, 0.1 mg ml^−1^ BSA, 0.01% Triton X-100, supplemented with 50 mM, 100 mM, or 150 mM NaCl) to minimize background ionic strength. Thus, the buffer series tested here corresponded to 70 mM, 120 mM, and 170 mM ions (at pH 7.5, about 10 mM HEPES is present as Na^+^HEPES^−^). By contrast, the original assay buffer was approximately 180 mM (40 mM from the NaCl and ∼140 mM from the sodium phosphate). After quenching reactions, NaCl was added to equalize the concentration across samples before adding DEAE resin. Quenching and subsequent steps were conducted using unmodified buffer.

For all assays, raw counts per minute (cpm) measurements were corrected for background signal by subtracting the average cpm of three mock (no enzyme) reactions processed in parallel. All activity assays were conducted in triplicate.

### Differential scanning fluorimetry

5 μM purified PWWP domain was incubated with 5× SYPRO Orange (Thermo Fisher Scientific) in assay buffer (20 mM Tris-HCl, pH 7.5, 150 mM NaCl, 5% glycerol). Samples were heated from 10 °C to 95 °C (10 s at each 0.5 °C step) using a CFX Connect qPCR (Bio-Rad). Melting temperatures were calculated with DSFWorld^61^ using model 1 in the “by sigmoid fitting” option. Assays were conducted in triplicate.

### Electrophoretic mobility shift assay

Purified PWWP domain at varying concentrations was incubated with 50 nM Cy3-labeled 30 bp DNA probe in assay buffer (20 mM HEPES, pH 7.5, 1 mM EDTA, 1 mg ml^−1^ BSA, 8% glycerol) for 20 min at room temperature. 5 μl of each binding reaction was loaded directly onto a pre-run 6% acrylamide DNA retardation gel (Thermo Fisher Scientific) and electrophoresed at ≤20 V cm^−1^ at 4 °C. Gels were imaged using a Sapphire Biomolecular Imager (Azure Biosystems).

### Fluorescence polarization assay

Purified PWWP domain was diluted to 8 μM in assay buffer (20 mM HEPES, pH 7.5, 1 mM EDTA, 0.5 mg ml^−1^ BSA, 8% glycerol) containing 20 nM Cy3-labeled 30 bp DNA probe (same as for EMSA). This was aliquoted in triplicate into a black 384-well plate (Corning), followed by 2-fold serial dilution in assay buffer containing 20 nM probe. The final volume in each well was 20 μl. Assay buffer was filtered before use, and stock solutions of probe and protein were centrifuged before use. The plate was incubated in the dark at room temperature for 1 h, and then read (1700 ms integration time) using a SpectraMax i3x with a rhodamine fluorescence polarization cartridge (Molecular Devices). Wells containing only assay buffer were used for background subtraction. Then, the G-factor was adjusted so that the polarization of wells containing assay buffer and 20 nM probe, but no protein, was set to a reference value of 27 mP. Curves were fit to the sigmoidal, 4PL model in GraphPad Prism.

### Peptide pulldown assay

1.5 nmol purified PWWP domain was incubated with 80 pmol biotinylated H3K36me0 or H3K36me2 peptide (same as in histone stimulation assays) in interaction buffer^5^ (50 mM Tris-HCl, pH 7.5 cold, 100 mM NaCl, 2 mM EDTA, 0.1% Triton X-100, fresh 0.5 mM DTT, fresh 0.2 mM phenylmethylsulfonyl fluoride, fresh 1× cOmplete EDTA-free protease inhibitor) overnight at 4 °C with rotation. To each sample, 10 μl Dynabeads MyOne T1 streptavidin beads were then added, and the mixtures were incubated at 4 °C with rotation for 4 h. The beads were washed 5 times with interaction buffer (10 min rotation at 4 °C per wash), and then boiled in loading buffer (95 °C, 5 min, 1000 rpm shaking). Eluted proteins were resolved by SDS-PAGE using a 10% tricine gel (Novex) and visualized by silver staining (Pierce).

### Generation of TKO ESC lines expressing ectopic Dnmt3a2

Approximately 3.6 × 10^4^ TKO ESCs were plated per well in a 12-well plate. The following day, cells were exchanged into medium without antibiotics and transfected with a 1:2 molar ratio of PBase:transposon plasmids, 1.1 μg total DNA per well, using FuGENE HD (3.5:1 ratio FuGENE:DNA). PBase was a gift from A. Meissner (Max Planck Institute). Cells were maintained in culture for 2 weeks, during which two rounds of FACS were used to isolate successfully transposed mCherry^+^ cells. 14 d after transfection, cells were MEF-depleted and harvested for analysis.

### Chromatin immunoprecipitation and sequencing (ChIP-seq)

19 × 10^6^ MEF-depleted parent TKO ESCs were crosslinked with 1% formaldehyde (Sigma-Aldrich) for 10 min at room temperature with rotation and then quenched with 125 mM glycine. Cells were pelleted, washed twice with cold PBS, snap frozen on dry ice, and stored at –80 °C. To perform ChIP, cells were first resuspended in 570 μl ChIP lysis buffer (50 mM Tris-HCl, pH 7.5, 1 mM EDTA, 1% SDS, 0.25% sodium deoxycholate (NaDOC), 1× Halt protease inhibitor) and incubated on ice for 10 min. Then, ChIP dilution buffer (50 mM Tris-HCl, pH 7.4, 150 mM NaCl, 0.01% SDS, 0.25% Triton X-100) was added to a final volume of 1.9 ml, and the lysate was sonicated using a Branson sonifier (0.7 s on, 1.3 s off, 5 min total on, 50% amplitude), followed by centrifugation to clarify lysates. The supernatant was diluted with ChIP dilution buffer to 5.7 ml, and a sample was reserved for ChIP input. The diluted lysate was incubated with antibody overnight at 4 °C with rotation: H3K4me3 (4 × 10^6^ cells, 2.5 μg anti-H3K4me3, Millipore, Cat#07-473, Lot#3394198), H3K36me2 (10 × 10^6^ cells, 1 μg anti-H3K36me2, Cell Signaling Technologies, Cat#2901, Lot#5). Dynabeads Protein G (Thermo Fisher Scientific) were then added and the samples were incubated for 2.5 h at 4 °C with rotation. The beads were isolated and washed twice with ChIP wash buffer A (10 mM Tris-HCl, pH 8.0, 150 mM NaCl, 1 mM EDTA, 0.1% SDS, 0.1% NaDOC, 1% Triton X-100), once with ChIP wash buffer B (10 mM Tris-HCl, pH 8.0, 250 mM LiCl, 0.5% NaDOC, 0.5% Triton X-100), and once with ChIP wash buffer C (10 mM Tris-HCl, pH 8.0, 50 mM NaCl, 1 mM EDTA). DNA was eluted by incubation (65 °C, 1 h, 1000 rpm shaking) with ChIP elution buffer (10 mM Tris-HCl, pH 8.0, 150 mM NaCl, 1 mM EDTA, 0.1% SDS). Each sample, ChIP or input (100 μL each), was then treated with RNase (50 μg, 37 °C, 30 min) and Proteinase K (25 μg, 63 °C, overnight), and purified by 2× SPRI. To construct libraries, 2.5 ng each sample was subjected to end-repair (End-It DNA End-Repair kit, Lucigen), A-tailing (Klenow fragment (3’-5’ exo–, New England Biolabs), ligation to barcoded adapters (KAPA), and PCR library amplification (KAPA library amplification primers; NEB Ultra 2× master mix), followed by double-sided SPRI size selection and a final 0.9× SPRI purification. Final libraries were analyzed by TapeStation (Agilent), pooled, and sequenced on a Novaseq SP flow cell with 2 × 50 bp reads.

### ChIP-seq data analysis

Reads were aligned to the mm10 genome (UCSC) using bwa-backtrack^62^ (v0.7.17-r1188). Samtools^63^ (v1.10) was used to convert sam files to bam files, which were then coordinate sorted and deduplicated using Picard (v2.26.9) (https://github.com/broadinstitute/picard). DeepTools bamCompare^64^ (v3.5.1) was used to create bigWig files for input-normalized ChIP signal, using 200 bp genomic bins and 1 kb smoothing (parameters: --extendReads --operation log2 --skipZeroOverZero -bs 200 --smoothLength 1000 --scaleFactorsMethod SES).

### Reduced representation bisulfite sequencing (RRBS)

RRBS was performed according to a published enhanced protocol^48^ with modifications. For each condition, genomic DNA was isolated from 2 × 10^5^ MEF-depleted ESCs using the QIAamp DNA Blood Mini kit. 80 ng genomic DNA was digested overnight with 150 units of MspI (New England Biolabs) at 37 °C. 0.5% unmethylated lambda phage DNA (Promega) was spiked in prior to digestion to enable downstream assessment of bisulfite conversion efficiency. The digest was purified by extraction with 25:24:1 phenol:chloroform:isoamyl alcohol and ethanol precipitation. Then, digested DNA was subjected to end-repair and A-tailing, and ligated to barcoded adapters (xGen Methyl UDI-UMI Adapters, Integrated DNA Technologies) using concentrated T4 ligase (New England Biolabs) at 16 °C overnight. Adapter-ligated DNA was purified by 1× SPRI, size-selected by gel extraction, bisulfite-converted (EZ DNA Methylation kit, Zymo; bisulfite conversion using the following thermocycler program: 55 cycles of 95 °C for 30 s, 50 °C for 15 min), and amplified using EpiMark hot start *Taq* DNA polymerase (New England Biolabs). Final libraries were analyzed by TapeStation, pooled, and sequenced using a Novaseq SP flow cell with 2 × 50 bp reads.

### RRBS data analysis

Sequencing reads were trimmed using Trim Galore (v0.6.7) (https://github.com/FelixKrueger/TrimGalore) with parameters --illumina --rrbs --paired --length 21. Reads 1 and 2 were swapped for trimming because the xGen Methyl UDI-UMI adapters used for library construction flip insert strandedness. Trimmed reads were aligned to the mm10 genome (UCSC) using Bismark^65^ (v0.23.1) (default bowtie2 alignment). Samtools^63^ (v1.10) was then used to sort and index bam files. MethylDackel (v0.6.1) (https://github.com/dpryan79/MethylDackel) was used to extract methylation at the CpG level, retaining only CpGs with at least 5× coverage (parameters: -d 5 --mergeContext --keepDupes). For subsequent analysis, we further filtered for only CpGs with at least 5× coverage across all samples using BedTools2 intersect^66^. The resulting bedGraph files were converted to bigWig format using bedGraphToBigWig (UCSC) and then processed using DeepTools multiBigwigSummary^64^ (v3.5.1) along with ChIP-seq bigWig files. A custom python script was used to remove CpGs/bins mapping to noncanonical chromosomes and perform additional processing (e.g., defining quartiles based on ChIP-seq signal).

### Statistics and replication

All statistical tests used were two-sided, and are described in the relevant figure legends. Tests were performed using SciPy (v1.2.0) or GraphPad Prism. Spearman correlations (Extended Data Fig. 2a) were calculated using Pandas (v1.0.1). Additional statistical details are provided in the figure legends or in the appropriate methods subsection. Unless otherwise noted in the figure legend, plots with error bars depict the mean ± SD of n = 3 replicates. Error bars are not shown in scatterplots (e.g., reporter silencing timecourses) where smaller than the data points themselves. Experiments were not randomized, and no statistical methods were used to select sample sizes. Methylation reporter assays (including sgRNA validation) and all biochemical experiments were each performed twice independently with similar results. Experiments involving next-generation sequencing (genotyping and genomics) were conducted once.

## References

1. Smith, Z. D. & Meissner, A. DNA methylation: roles in mammalian development. Nature Reviews Genetics 14, 204–20 (2013).

2. Schübeler, D. Function and information content of DNA methylation. Nature 517, 321–326 (2015).

3. Okano, M., Bell, D. W., Haber, D. A. & Li, E. DNA Methyltransferases Dnmt3a and Dnmt3b Are Essential for De Novo Methylation and Mammalian Development. Cell 99, 247–257 (1999).

4. Tatton-Brown, K. et al. Mutations in the DNA methyltransferase gene DNMT3A cause an overgrowth syndrome with intellectual disability. Nature Genetics 46, ng.2917 (2014).

5. Heyn, P. et al. Gain-of-function DNMT3A mutations cause microcephalic dwarfism and hypermethylation of Polycomb-regulated regions. Nature Genetics 51, 1–10 (2019).

6. Jaiswal, S. & Ebert, B. L. Clonal hematopoiesis in human aging and disease. Science 366, eaan4673 (2019).

7. Ley, T. J. et al. DNMT3A Mutations in Acute Myeloid Leukemia. The New England Journal of Medicine 363, 2424–2433 (2010).

8. Brunetti, L., Gundry, M. C. & Goodell, M. A. DNMT3A in Leukemia. Cold Spring Harbor Perspectives in Medicine 7, a030320 (2017).

9. Shlush, L. I. et al. Identification of pre-leukaemic haematopoietic stem cells in acute leukaemia. Nature 506, 328 (2014).

10. Jeong, M. et al. Loss of Dnmt3a Immortalizes Hematopoietic Stem Cells In Vivo. Cell Reports 23, 1–10 (2018).

11. Mayle, A. et al. Dnmt3a loss predisposes murine hematopoietic stem cells to malignant transformation. Blood 125, 629–38 (2015).

12. Suetake, I., Shinozaki, F., Miyagawa, J., Takeshima, H. & Tajima, S. DNMT3L Stimulates the DNA Methylation Activity of Dnmt3a and Dnmt3b through a Direct Interaction. Journal of Biological Chemistry 279, 27816–27823 (2004).

13. Holz-Schietinger, C., Matje, D. M., Harrison, M. F. & Reich, N. O. Oligomerization of DNMT3A Controls the Mechanism of de Novo DNA Methylation. Journal of Biological Chemistry 286, 41479–41488 (2011).

14. Xu, T.-H. et al. Structure of nucleosome-bound DNA methyltransferases DNMT3A and DNMT3B. Nature 1–5 (2020) doi:10.1038/s41586-020-2747-1.

15. Duymich, C. E., Charlet, J., Yang, X., Jones, P. A. & Liang, G. DNMT3B isoforms without catalytic activity stimulate gene body methylation as accessory proteins in somatic cells. Nature Communications 7, 11453 (2016).

16. Nguyen, T.-V. et al. The R882H DNMT3A hot spot mutation stabilizes the formation of large DNMT3A oligomers with low DNA methyltransferase activity. J Biol Chem 294, 16966–16977 (2019).

17. Guo, X. et al. Structural insight into autoinhibition and histone H3-induced activation of DNMT3A. Nature 517, 640 (2015).

18. Weinberg, D. N. et al. The histone mark H3K36me2 recruits DNMT3A and shapes the intergenic DNA methylation landscape. Nature 573, 281–286 (2019).

19. Xu, W. et al. DNMT3A reads and connects histone H3K36me2 to DNA methylation. Protein Cell 11, 150–154 (2020).

20. Bröhm, A. et al. Methylation of recombinant mononucleosomes by DNMT3A demonstrates efficient linker DNA methylation and a role of H3K36me3. Commun Biology 5, 192 (2022).

21. Zhang, Z.-M. et al. Structural basis for DNMT3A-mediated de novo DNA methylation. Nature 554, 387 (2018).

22. Noh, K.-M. et al. Engineering of a Histone-Recognition Domain in Dnmt3a Alters the Epigenetic Landscape and Phenotypic Features of Mouse ESCs. Molecular Cell 59, 89–103 (2015).

23. Wu, H. et al. Structural and Histone Binding Ability Characterizations of Human PWWP Domains. Plos One 6, e18919 (2011).

24. Huang, Y.-H. et al. Systematic profiling of DNMT3A variants reveals protein instability mediated by the DCAF8 E3 ubiquitin ligase adaptor. Cancer Discov 12, candisc.0560.2021 (2021).

25. Vinyard, M. E. et al. CRISPR-suppressor scanning reveals a nonenzymatic role of LSD1 in AML. Nature Chemical Biology 15, 529–539 (2019).

26. Gosavi, P. M. et al. Profiling the Landscape of Drug Resistance Mutations in Neosubstrates to Molecular Glue Degraders. Acs Central Sci (2021) doi:10.1021/acscentsci.1c01603.

27. Shen, C. et al. NSD3-Short Is an Adaptor Protein that Couples BRD4 to the CHD8 Chromatin Remodeler. Molecular Cell 60, 847–859 (2015).

28. Shi, J. et al. Discovery of cancer drug targets by CRISPR-Cas9 screening of protein domains. Nature Biotechnology 33, nbt.3235 (2015).

29. Sher, F. et al. Rational targeting of a NuRD subcomplex guided by comprehensive in situ mutagenesis. Nature Genetics 51, 1149–1159 (2019).

30. Wang, T. et al. Identification and characterization of essential genes in the human genome. Science (New York, N.Y.) 350, 1096–101 (2015).

31. Komor, A. C., Kim, Y. B., Packer, M. S., Zuris, J. A. & Liu, D. R. Programmable editing of a target base in genomic DNA without double-stranded DNA cleavage. Nature 533, 420 (2016).

32. Hanna, R. E. et al. Massively parallel assessment of human variants with base editor screens. Cell 184, 1064-1080.e20 (2021).

33. Cuella-Martin, R. et al. Functional interrogation of DNA damage response variants with base editing screens. Cell 184, 1081-1097.e19 (2021).

34. Sangree, A. K. et al. Benchmarking of SpCas9 variants enables deeper base editor screens of BRCA1 and BCL2. Nat Commun 13, 1318 (2022).

35. Xu, P. et al. Genome-wide interrogation of gene functions through base editor screens empowered by barcoded sgRNAs. Nat Biotechnol 1–11 (2021) doi:10.1038/s41587-021-00944-1.

36. Bintu, L. et al. Dynamics of epigenetic regulation at the single-cell level. Science 351, 720– 724 (2016).

37. Barretina, J. et al. The Cancer Cell Line Encyclopedia enables predictive modelling of anticancer drug sensitivity. Nature 483, 603 (2012).

38. Reither, S., Li, F., Gowher, H. & Jeltsch, A. Catalytic Mechanism of DNA-(cytosine-C5)-methyltransferases Revisited: Covalent Intermediate Formation is not Essential for Methyl Group Transfer by the Murine Dnmt3a Enzyme. Journal of Molecular Biology 329, 675–684 (2003).

39. Holz-Schietinger, C., Matje, D. M. & Reich, N. O. Mutations in DNA Methyltransferase (DNMT3A) Observed in Acute Myeloid Leukemia Patients Disrupt Processive Methylation. Journal of Biological Chemistry 287, 30941–30951 (2012).

40. Tate, J. G. et al. COSMIC: the Catalogue Of Somatic Mutations In Cancer. Nucleic Acids Res 47, D941–D947 (2019).

41. Li, B.-Z. et al. Histone tails regulate DNA methylation by allosterically activating de novo methyltransferase. Cell Research 21, 1172 (2011).

42. Sievers, Q. L., Gasser, J. A., Cowley, G. S., Fischer, E. S. & Ebert, B. L. Genome-wide screen identifies cullin-RING ligase machinery required for lenalidomide-dependent CRL4CRBN activity. Blood 132, 1293–1303 (2018).

43. Tatton-Brown, K. et al. The Tatton-Brown-Rahman Syndrome: A clinical study of 55 individuals with de novo constitutive DNMT3A variants. Wellcome Open Res 3, 46 (2018).

44. Dukatz, M. et al. H3K36me2/3 binding and DNA binding of the DNA methyltransferase DNMT3A PWWP domain both contribute to its chromatin interaction. J Mol Biol 431, 5063–5074 (2019).

45. Purdy, M. M., Holz-Schietinger, C. & Reich, N. O. Identification of a second DNA binding site in human DNA methyltransferase 3A by substrate inhibition and domain deletion. Arch Biochem Biophys 498, 13–22 (2010).

46. Suetake, I. et al. Characterization of DNA-binding activity in the N-terminal domain of the DNA methyltransferase Dnmt3a. Biochemical Journal 437, 141–148 (2011).

47. Haggerty, C. et al. Dnmt1 has de novo activity targeted to transposable elements. Nat Struct Mol Biol 28, 1–10 (2021).

48. Garrett-Bakelman, F. E. et al. Enhanced Reduced Representation Bisulfite Sequencing for Assessment of DNA Methylation at Base Pair Resolution. J Vis Exp Jove 52246 (2015) doi:10.3791/52246.

49. Gu, H. et al. Preparation of reduced representation bisulfite sequencing libraries for genome-scale DNA methylation profiling. Nat Protoc 6, 468–481 (2011).

50. Weinberg, D. N. et al. Two competing mechanisms of DNMT3A recruitment regulate the dynamics of de novo DNA methylation at PRC1-targeted CpG islands. Nat Genet 53, 794–800 (2021).

51. Tycko, J. et al. High-Throughput Discovery and Characterization of Human Transcriptional Effectors. Cell 183, 2020-2035.e16 (2020).

52. Clement, K. et al. CRISPResso2 provides accurate and rapid genome editing sequence analysis. Nat Biotechnol 37, 224–226 (2019).

53. Li, L.-C. & Dahiya, R. MethPrimer: designing primers for methylation PCRs. Bioinformatics 18, 1427–1431 (2002).

54. Joung, J. et al. Genome-scale CRISPR-Cas9 knockout and transcriptional activation screening. Nature Protocols 12, 828 (2017).

55. Canver, M. C. et al. Integrated design, execution, and analysis of arrayed and pooled CRISPR genome-editing experiments. Nature Protocols 13, 946 (2018).

56. Kamburov, A. et al. Comprehensive assessment of cancer missense mutation clustering in protein structures. Proc National Acad Sci 112, E5486–E5495 (2015).

57. Eddy, S. R. Accelerated Profile HMM Searches. Plos Comput Biol 7, e1002195 (2011).

58. Suzek, B. E. et al. UniRef clusters: a comprehensive and scalable alternative for improving sequence similarity searches. Bioinformatics 31, 926–932 (2015).

59. El-Gebali, S. et al. The Pfam protein families database in 2019. Nucleic Acids Res 47, D427–D432 (2019).

60. Sievers, F. et al. Fast, scalable generation of high-quality protein multiple sequence alignments using Clustal Omega. Mol Syst Biol 7, 539–539 (2011).

61. Wu, T. et al. Three Essential Resources to Improve Differential Scanning Fluorimetry (DSF) Experiments. Biorxiv 2020.03.22.002543 (2020) doi:10.1101/2020.03.22.002543.

62. Li, H. & Durbin, R. Fast and accurate short read alignment with Burrows–Wheeler transform. Bioinformatics 25, 1754–1760 (2009).

63. Danecek, P. et al. Twelve years of SAMtools and BCFtools. Gigascience 10, giab008 (2021).

64. Ramírez, F. et al. deepTools2: a next generation web server for deep-sequencing data analysis. Nucleic Acids Res 44, W160–W165 (2016).

65. Krueger, F. & Andrews, S. R. Bismark: a flexible aligner and methylation caller for Bisulfite-Seq applications. Bioinformatics 27, 1571–1572 (2011).

66. Quinlan, A. R. & Hall, I. M. BEDTools: a flexible suite of utilities for comparing genomic features. Bioinformatics 26, 841–842 (2010).

